# ADAR1 interaction with Z-RNA promotes editing of endogenous double-stranded RNA and prevents MDA5-dependent immune activation

**DOI:** 10.1101/2020.12.04.411702

**Authors:** Richard de Reuver, Evelien Dierick, Bartosz Wiernicki, Katrien Staes, Leen Seys, Ellen De Meester, Tuur Muyldermans, Alexander Botzki, Bart Lambrecht, Filip Van Nieuwerburgh, Peter Vandenabeele, Jonathan Maelfait

## Abstract

Loss-of-function of ADAR1 causes the severe autoinflammatory disease Aicardi-Goutières Syndrome (AGS). ADAR1 converts adenosines into inosines within double-stranded (ds) RNA. This process called A-to-I editing masks self-dsRNA from detection by the antiviral dsRNA sensor MDA5. ADAR1 binds to dsRNA in both the canonical A-form and in the poorly defined Z-conformation (Z-RNA). Mutations in the Z-RNA binding Zα-domain of ADAR1 are common in AGS patients. How loss of ADAR1/Z-RNA interaction contributes to disease development is unknown. Using ADAR1 Zα-domain mutant human cells and knock-in mice, we demonstrate that abrogated binding of ADAR1 to Z-RNA leads to reduced A-to-I editing of dsRNA structures formed by pairing of inversely oriented SINEs. As a result, ADAR1 Zα-domain mutant human cells and transgenic mice develop a spontaneous MDA5-dependent immune response. This shows that the interaction between ADAR1 and Z-RNA restricts sensing of self-dsRNA and prevents AGS development.

## Introduction

Aicardi-Goutières Syndrome (AGS) is a severe autoinflammatory disease characterised by chronic type-I interferon (IFN) production (Rodero and Crow, 2016). AGS results from mutations in one of seven genes including *ADAR*, which codes for Adenosine Deaminase Acting on dsRNA-1 (ADAR1) (Crow et al., 2015; Rice et al., 2012). ADAR1 converts adenosine (A) residues into inosines (I) within double-stranded (ds) RNA. This process called A-to-I editing marks endogenous dsRNA as self and masks these molecules from detection by the cytosolic nucleic acid sensor MDA5. Mutations that disrupt the activity of ADAR1 such as in AGS trigger a deleterious MDA5-mediated type-I IFN response.

ADAR1 mainly edits dsRNA structures formed by base pairing of two adjacent and inversely oriented short interspersed nuclear elements (SINEs). These SINEs include the human *Alu* elements and mouse B1 or B2 repeats, which are abundantly present in introns and 3’ untranslated regions (3’UTRs) of messenger RNAs (mRNAs) (Athanasiadis et al., 2004; Blow et al., 2004; Kim et al., 2004; Levanon et al., 2004; Neeman et al., 2006). A-to-I editing of endogenous SINE-derived duplex RNA thus forms an important barrier to prevent spontaneous MDA5 activation (Ahmad et al., 2018; Chung et al., 2018). This is supported by the analysis of *Adar* knockout or editing-deficient mice, which develop an MDA5-mediated type I IFN response (Bajad et al., 2020; Liddicoat et al., 2015; Mannion et al., 2014; Pestal et al., 2015).

*ADAR* encodes two isoforms: a constitutively expressed short transcript (p110) and a long isoform (p150) that is transcribed from an IFN-inducible promotor. ADAR1 binds to the canonical A-conformation of dsRNA (A-RNA) while the larger p150 isoform contains an N-terminal Zα-domain, which specifically interacts with nucleic acids that have adopted the unusual Z-conformation, including Z-RNA (Placido et al., 2007). Recent evidence shows that both viral and endogenous Z-RNA sequences bind to the tandem Zα-domains of the nucleic acid sensor ZBP1. This then triggers ZBP1 activation and contributes to cell-intrinsic antiviral immunity or – when aberrantly activated – to the development of inflammatory disease (Devos et al., 2020; Jiao et al., 2020; Kesavardhana et al., 2017; Kesavardhana et al., 2020; Maelfait et al., 2017; Sridharan et al., 2017; Wang et al., 2020; Zhang et al., 2020). These findings revealed a role for the recognition of Z-RNA by Zα-domains of ZBP1 in inducing an immune response. How Z-RNA relates to ADAR1 function and whether the Zα-domain in ADAR1 is functionally important for its immunosuppressive effects remains unknown.

Mice that only express ADAR1-p110 and that are deficient for the Zα-domain containing p150 isoform phenocopy full *Adar*-deficient animals and develop an MDA5-dependent type-I IFN response (Pestal et al., 2015; Ward et al., 2011). Interestingly, more than half of *ADAR* AGS patients contain compound heterozygous mutations with one allele carrying the p.Pro193Ala mutation in the Zα-domain of ADAR1-p150 combined with another dysfunctional *ADAR* allele (Rice et al., 2012; Rice et al., 2017). Together, this strongly suggests that the A-to-I editing activity of the ADAR1-p150 isoform is the primary mechanism by which MDA5 evades recognition of endogenous dsRNA and that the interaction of ADAR1 with Z-RNA contributes to this inhibitory activity. However, experimental evidence that supports this hypothesis is currently lacking.

We generated human cell lines and *Adar* knock-in mice expressing an ADAR1 protein that is unable to interact with Z-RNA, while leaving the A-form dsRNA binding motifs and deaminase activity of ADAR1 intact. We found that Z-RNA binding to ADAR1 is required to enable efficient A-to-I editing of duplex RNA formed by oppositely oriented human *Alu* or mouse B1 or B2 SINEs. Disruption of the Zα-domain of ADAR1 led to the accumulation of dsRNA and triggered an MDA5-mediated type-I IFN response. Homozygous *Adar* Zα-domain mutant mice developed a systemic antiviral immune response reminiscent of AGS patients, showed mild signs of spontaneous adaptive immune activation and were more resistant to viral infection, but did not develop autoinflammatory disease. We propose that the Zα-domain of ADAR1 widens its editing repertoire by interacting with Z-RNA structures in duplex RNA substrates, which may otherwise not be accessible to the A-RNA binding dsRBMs. Binding of ADAR1 to Z-RNA therefore contributes to the discrimination between self and non-self and is important in preventing genetic disease such as AGS.

## Results

### ADAR1 Zα-domain mutation activates MDA5 in human cells

The *ADAR* p.Pro193Ala (c.577C>G) missense mutation located in the Z-form nucleic acid binding Zα-domain of ADAR1 has an allele frequency of ∼0.2%. p.Pro193Ala occurs in the compound heterozygous state with a dysfunctional *ADAR* allele in around half of *ADAR* AGS cases (Fig. 1A) (Rice et al., 2017). Proline 193 is located in the β-loop that connects the Z-form nucleic acid recognition α3-helix with two C-terminal antiparallel β2- and β3-sheets in the Zα-domain of ADAR1 (Placido et al., 2007; Schwartz et al., 1999). Proline mutations often result in structural irregularities and may cause Zα-domain misfolding. To specifically address the impact of disrupted binding of ADAR1 to Z-RNA, we decided to introduce N173A/Y177A mutations in the Zα-domain of ADAR1, which fully abrogate binding to Z-form nucleic acids without altering overall Zα-domain structure (Feng et al., 2011; Schade et al., 1999). Using CRISPR/Cas9-mediated homology directed repair we generated four HEK293 clones with Zα-domain mutant *ADAR* (*ADAR*^Zαmut^) alleles (Figure S1A and S1B). All alleles carried the Y177A mutation, however, we detected at least one intact N173 allele in the four clones (Figure S1A), possibly due to incomplete targeting of the hypo-triploid HEK293 genome (Bylund et al., 2004). Sole Y177A mutation reduces the affinity of the Zα-domain for Z-form nucleic acids by 30-fold (Schade et al., 1999). We reasoned that the remaining Y177A-only protein expressed by this locus would still be compromised in its ability to bind to Z-RNA and would not interfere with the interpretation of our results. As controls, we retained two wild type (*ADAR*^WT^) clones and generated an ADAR1 deficient (*ADAR*^KO^) clone (Figure S1A and S1B). ADAR1 p110 or p150 protein expression was not affected by introducing N173A/Y177A mutations in the Zα-domain of ADAR1 (Figure S1B). Loss of ADAR1 in HEK293T cells induced an MDA5-dependent interferon response and leads to a spontaneous activation of the dsRNA sensor and translational inhibitor Protein Kinase R (PKR) (Ahmad et al., 2018; Chung et al., 2018; Pestal et al., 2015). Ectopic expression of MDA5 in *ADAR*^KO^ but not in *ADAR*^WT^ or parental cells induced strong activation of an IFN-β reporter (Figure 1B) confirming these results. MDA5 expression in *ADAR*^Zαmut^ cells similarly resulted in an elevated IFN-β promotor response upon MDA5 expression albeit less robustly than in ADAR1 deficient cells (Figure 1B). The increased IFN response was specific for MDA5 as overexpression of TRIF, an adaptor molecule for TLR3/TLR4-mediated type-I IFN induction, induced equal responses regardless of the *ADAR* genotype (Figure S1C). This suggests that endogenous agonists of MDA5 accumulate in *ADAR*^Zαmut^ cells. Indeed, co-expression of the viral dsRNA-binding proteins NS1 from Influenza A or E3L from Vaccinia virus, but not the dsRNA-binding mutant proteins prevented the MDA5-induced IFN response (Figure 1C).

**Figure 1.**
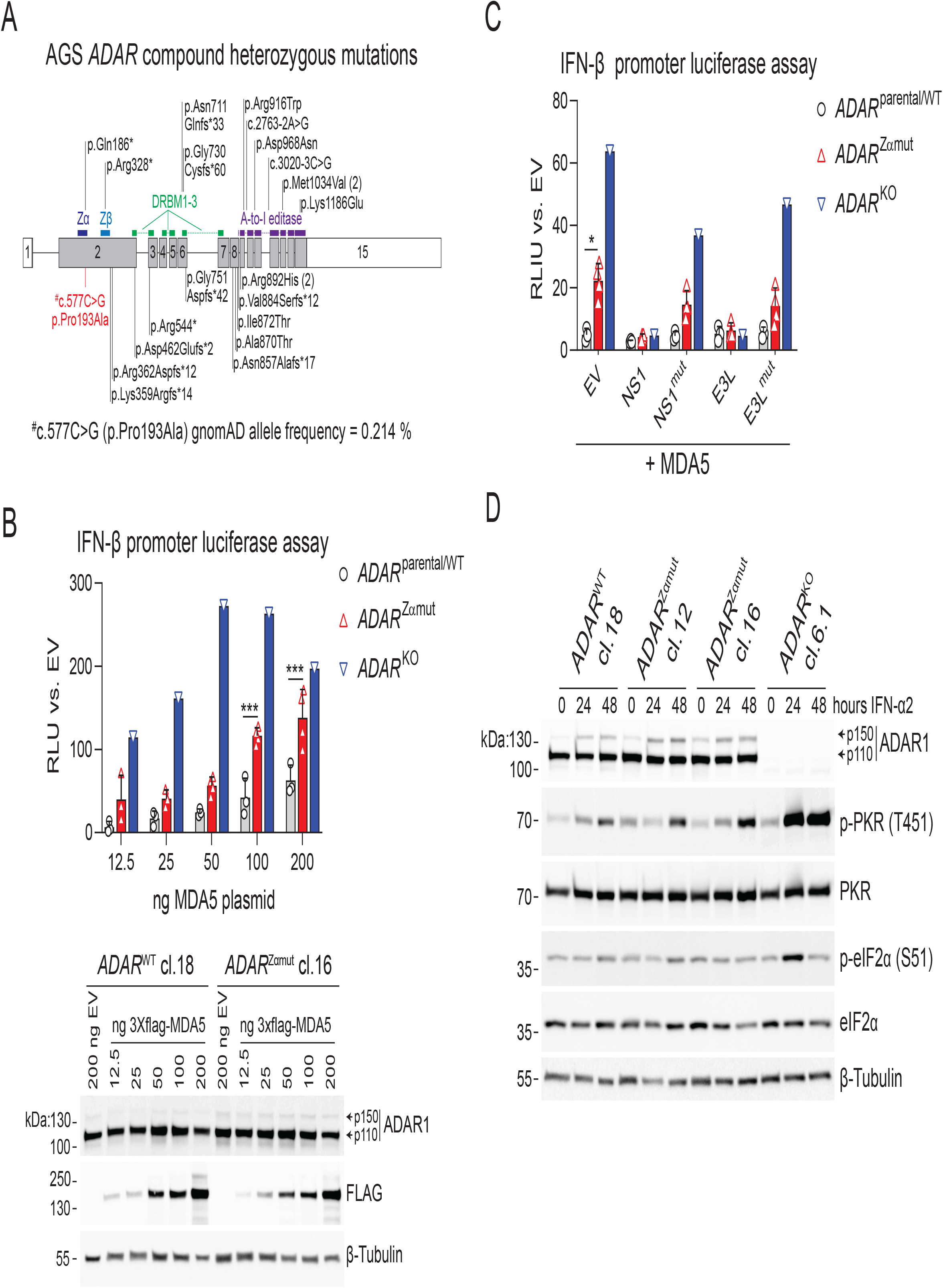
ADAR1 Zα-domain mutation activates MDA5 in human cells. **(A)** Schematic overview of the *ADAR* gene encoding for ADAR1. Exons are indicated as boxes and introns as lines. The coding sequence is in grey and the 5’UTR and 3’UTR is in white. Protein domains including the Z-RNA binding Zα-domain, a Zβ domain, three dsRNA-binding motifs (DRBMs) and the A-to-I editase domain are indicated as colored bars. Compound heterozygous ADAR mutation that occur in AGS patients are shown. The Zα-domain mutation c.577C>G (p.Pro193Ala) is indicated in red. *ADAR* nonsense mutations or missense mutations that are predicted to abrogate the ADAR1 A-to-I editing activity of the other allele are in black. **(B)** Parental HEK293 cells or *ADAR* wild type clones (*ADAR*^parental/WT^), Zα-domain mutant (*ADAR*^Zαmut^) or ADAR1-deficient (*ADAR*^Zαmut^) HEK293 clones were transfected with 50 ng IFN-β promoter firefly luciferase reporter and 20 ng *Renilla* luciferase reporter plasmids, together with increasing amounts of FLAG-tagged human MDA5. Luciferase activity was measured after 24 h, and the relative luciferase units (RLU) were calculated by determining the ratio of firefly and *Renilla* luciferase units. RLU was set to 1 for cells transfected with 200 ng empty vector. Protein expression of ADAR1 and FLAG-tagged MDA5 in a *ADAR*^WT^ and *ADAR*^Zαmut^ clone was verified by Western blot (bottom panel). Arrows indicate the ADAR1 p110 and p150 isoform. ***, P < 0.001 by 2-way ANOVA. **(C)** Cells of the indicated genotype with IFN-β promoter firefly luciferase and *Renilla* luciferase reporter plasmids and 100 ng FLAG-tagged human MDA5 together with 100 ng of wild type NS1 or NS1^R38A/K41A^ (NS1^mut^) from influenza A or wild type E3L or E3L^N44A/Y48A/K167A^ (E3L^mut^). Luciferase activity was analysed as in (B). *, P < 0.05 by unpaired t-test. Bars in (B) and (C) represent mean normalised F-luc expression and symbols represent individual clones. **(D)** HEK293 clones of the indicated genotypes were left untreated or stimulated for 24 or 48 h with 1000 U/ml IFNα2. Protein expression of ADAR1, threonine 451 (T451) phosphorylated PKR (p-PKR), total PKR, serine 51 (S51) phosphorylated eIF2α (p-eIF2α) and total eIF2α was analysed by Western blotting. Arrows indicate the ADAR1 p110 and p150 isoform. Data in (B), (C) and (D) are representative of at least two independent experiments. See also Figure S1.

Next, we tested whether a spontaneous PKR response occurred in *ADAR*^Zαmut^ HEK293 cells. As reported previously (Chung et al., 2018), IFN-α2 treatment of *ADAR*^KO^ cells induced strong activation of the kinase activity of PKR as shown by PKR autophosphorylation on Threonine 451 and Serine 51 phosphorylation of the of alpha subunit of its substrate eukaryotic initiation factor 2 (eIF2α) at 24 and/or 48 hours post stimulation (Figures 1D and S1D). In contrast, Zα-domain mutation led to a marginal increase in activation of PKR compared to cells expressing wild type ADAR1 only at 48 hours after IFN-α2 treatment (Figures 1D and S1D). Together, these data indicate that the interaction between the Zα-domain of ADAR1 and Z-RNA suppresses the development of a spontaneous MDA5-mediated type-I IFN response but does not affect PKR activation.

### *Adar* Zα-domain mutant mice develop a spontaneous type-I IFN response

To study the impact of ADAR1 Zα-domain mutation *in vivo*, we generated *Adar* knock-in mice expressing an ADAR1 protein in which we mutated the Zα-domain residues N175 and Y179 into alanines, orthologous to the human N173A/Y177A variant in the HEK293 cells. As complete ADAR1 and ADAR1-p150 knockout mice die embryonically (Hartner et al., 2004; Wang et al., 2004; Ward et al., 2011), we decided to create conditional ADAR1 Zα-domain knock-in mice by cloning an inverted mutant Zα-domain containing exon 2 downstream of the wild type sequence. Both the wild type and mutant exons were flanked by LoxP and Lox2272 sites in such a way that Cre-mediated excision of the wild type exon and inversion of the Zα-domain mutant exon creates a Zα-domain mutant *Adar* allele (*Adar*^Zα^) (Figure S2A). First, we crossed heterozygous male conditional Zα-domain mutant mice (*Adar*^fl-Zα/+)^ to female Sox2-Cre mice enabling epiblast-specific recombination of the *Adar*^fl-Zα^ allele into a Zα-domain mutant *Adar*^Zα^ allele (Hayashi et al., 2003). The resulting heterozygous *Adar*^Zα/+^ offspring were interbred to generate homozygous *Adar*^Zα/Zα^ mice. Sanger sequencing confirmed complete Cre-mediated recombination (Figure S2B). In contrast to *Adar*^-/-^ and *Adar-p150*^-/-^ mice, *Adar*^Zα/Zα^ animals were born at Mendelian ratios, had no gross defects and bred normally (Figure S2C). Importantly, Zα-domain mutation did not affect ADAR1 p110 or p150 protein expression in bone marrow-derived macrophages (BMDMs) or primary lung fibroblasts in resting or type-I IFN-stimulated cells (Figure 2A and S2D). ADAR1 p110 expression is mainly nuclear while p150 shuttles between the nucleus and the cytosol. We confirmed that ADAR1 localisation did not differ in *Adar*^Zα/Zα^ or control fibroblasts (Figure S2D).

**Figure 2.**
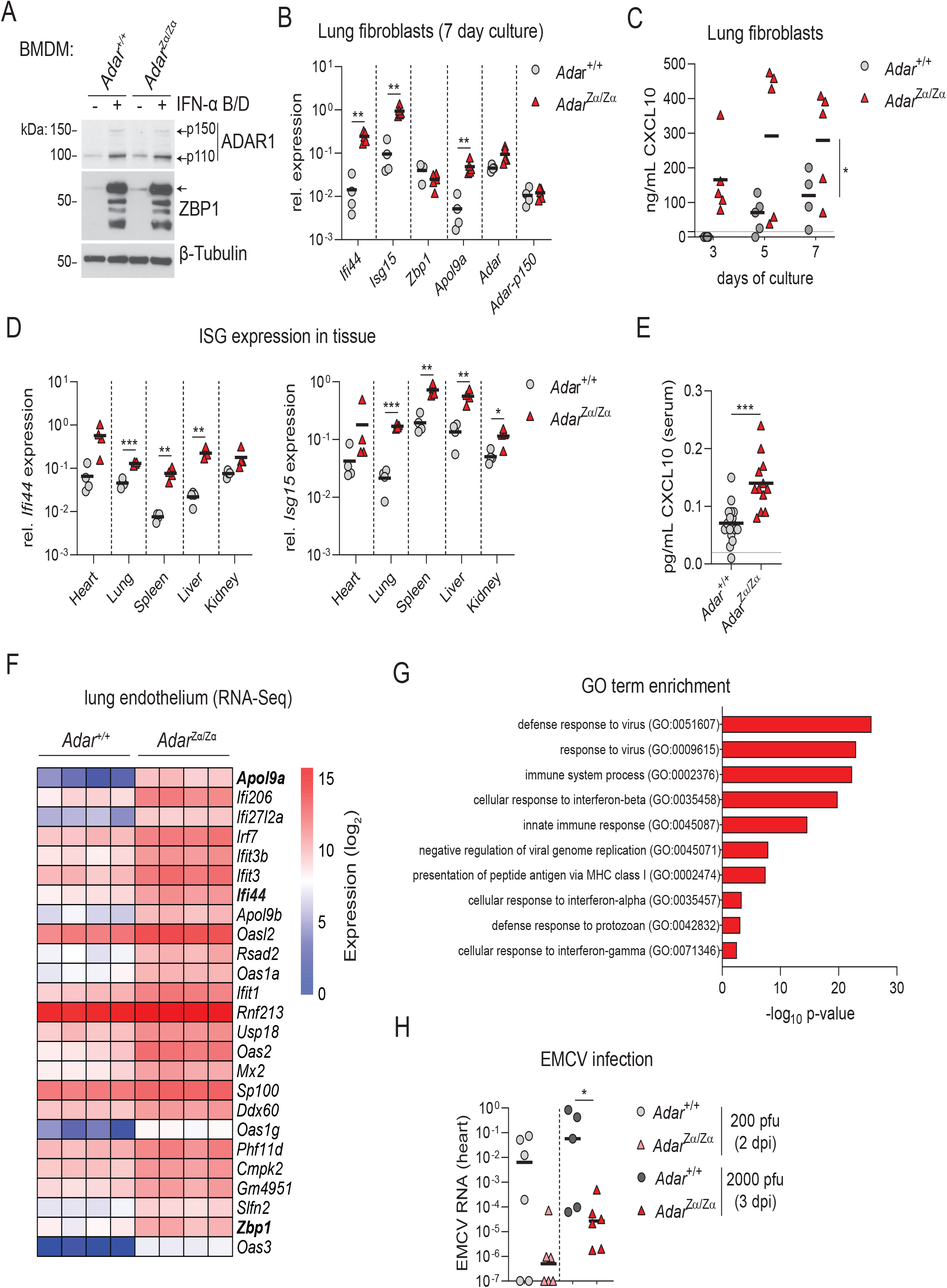
*Adar* Zα-domain mutant mice develop a spontaneous type-I IFN response. **(A)** Bone-marrow derived macrophages (BMDMs) from wild type (*Adar*^+/+^) and ADAR1 Zα-domain mutant (Adar^Zα/Zα^) mice were left untreated or stimulated for 18 h with 200 U/ml IFN-α B/D. Protein expression of ADAR1 (isoform p110 and p150) and ZBP1 was analysed by Western blot. **(B)** Primary lung fibroblasts from *Adar*^+/+^ and *Adar*^Zα/Zα^ mice were cultured without medium change for 7 days. (B) RT-qPCR analysis of the indicated ISGs at day 7 of the culture. **, P < 0.01 by unpaired t-test. **(C)** CXCL10 protein levels in the cell culture supernatant from (B) were measured on day 3, 5 and 7 by ELISA. *, P < 0.05 by 2-way ANOVA. Each data point in (B) and (C) represents a fibroblast culture from an individual mouse. Lines indicate the mean. **(D)** RT-qPCR analysis of *Ifi44* and *Isg15* in the indicated organs of 8-10 week old *Adar*^+/+^ and *Adar*^Zα/Zα^ mice. *, P < 0.05, **, P < 0.01, ***, P < 0.001 by unpaired t-test. **(E)** Analysis of CXCL10 protein levels in serum of 16 week old *Adar*^+/+^ and *Adar*^Zα/Zα^ mice by Bio-Plex assay. ***, P < 0.001 by Mann-Whitney U test. Each data point in (D) and (E) represents an individual mouse. Lines indicate the mean. **(F)** Log2 normalised expression levels of top 25 (total: n = 152, see Supplementary Table 1) differentially expressed genes (DEGs) identified by total stranded RNA-seq data from primary lung endothelial cells of *Adar*^+/+^ (n = 4) and *Adar*^Zα/Zα^ (n = 4) mice. **(G)** Gene ontology (GO) enrichment analysis of all 152 DEGs identified in (F) by the functional annotation tool of DAVID with Bonferroni correction. **(H)** *Adar*^+/+^ and *Adar*^Zα/Zα^ mice were infected with 200 or 2000 plaque forming units (pfu) of EMCV. At 2 or 3 days post-infection (dpi) respectively, viral titres were determined in the hearts by RT-qPCR of EMCV RNA. Line represent the mean. Symbols represent individual mice. *, P < 0.05 by Mann–Whitney U test. Data in (A) to (D) and (H) are representative of at least 2 independent experiments. See also Figure S2 and S3.

As the Zα-domain of ADAR1 is required to suppress MDA5-mediated type-I IFN inducation in human cells (see Figure 1B), we analysed mRNA expression of the interferon-stimulated genes (ISGs) *Ifi44, Isg15* and *Zbp1* in *Adar*^Zα/Zα^ cells. While overnight BMDM or lung fibroblast cultures did not develop a type-I IFN response (Figures S2E and S2F), we detected a strong type-I IFN gene signature when *Adar*^Zα/Zα^ lung fibroblasts were cultured for 1 week (Figure 2B). This delayed response most likely depended on a feedforward loop induced by the autocrine secretion of type-I IFNs in the cell culture medium. Indeed, *Adar*^Zα/Zα^ cells chronically produced increased levels of CXCL10 protein, a sensitive surrogate marker for type-I IFNs (Figure 2C). Analysis of *Ifi44* and *Isg15* expression across several tissues of 8-10 week old mice including heart, lung, spleen, liver and kidney also revealed a type-I IFN gene signature in *Adar*^Zα/Zα^ animals (Figure 2D). In line with this, we detected enhanced levels of CXCL10 protein in serum and increased protein expression of the ISG and antiviral nucleic acid sensor ZBP1 in spleen and lung tissue (Figures 2E and S2G) of *Adar*^Zα/Zα^ animals.

Similar to human cells, ADAR1 deficiency in mouse cells also leads to spontaneous PKR activation (see Figures 1D and S1D) and triggers an unfolded protein response (UPR) during mesenchymal-to-epithelial transition (George et al., 2016; Guallar et al., 2020). Phosphorylation of eIF2α by PKR halts initiation of general protein translation and promotes selective induction of ATF4-dependent genes including asparagine synthase (*Asns*) and CHOP (*Ddit3*) as part of the integrated stress response (Donnelly et al., 2013). To determine whether ADAR1 Zα-domain mutation triggers PKR/ATF4 activation and/or induces an UPR, we measured mRNA expression of the ATF4-target genes *Asns* and *Ddit3* and general UPR-induced transcripts including *BiP* (or *Hspa5*) and *Xbp1* in the unspliced (*Xbp1u*) and spliced (*Xbp1s*) forms in the lungs of *Adar*^Zα/Zα^ mice (Figure S2H). In contrast to the robust type-I IFN signature (Figure 2D), we did not find evidence for PKR or UPR activation.

We then asked whether various cell types were differentially affected by loss of ADAR1/Z-RNA interactions. To this end, we sorted CD45-positive leukocytes including alveolar macrophages, neutrophils, conventional type 1 and type 2 dendritic cells (cDC1 and cDC2) and CD45-negative endothelial and epithelial cells from lungs (Figures S3A and S3B) as well as B cells, T cells and NK cells from spleens (Figure S3C) of *Adar*^Zα/Zα^ mice and control littermates. Analysis of mRNA expression of several of ISGs showed that the Zα-domain of ADAR1 is essential to suppress a spontaneous type-I IFN response in all cell types, although the magnitude of the response was different (Figure S3D and S3E). Neutrophils, lung epithelium and lung endothelial cells displayed a particularly robust type-I IFN signature (Figure S3D). It remains to be determined whether all these cell types act as source of type-I IFNs or that they responded to elevated circulating type-I IFNs in a non-cell autonomous manner. RNA-Seq of FACS-purified lung endothelial cells confirmed that loss of ADAR1 interaction with Z-RNA triggered a spontaneous type-I IFN response. All top 25 upregulated differentially expressed genes were ISGs (Figure 2F) and GO enrichment analysis did not reveal any signs of activation of other immune pathways (Figure 2G). Finally, to determine whether the spontaneous antiviral immune response of *Adar*^Zα/Zα^ mice is protective against viruses, we infected Zα-domain mutant mice and control littermates with two doses of encephalomyocarditis virus (EMCV), a positive stranded RNA virus that is recognised by MDA5 (Gitlin et al., 2006). In line with their enhanced antiviral state, we detected highly reduced copies of EMCV RNA in the hearts of *Adar*^Zα/Zα^ mice compared to control animals at 2 and 3 days post infection (Figure 2H).

We conclude that disruption of the interaction between ADAR1 and Z-RNA in *Adar*^Zα/Zα^ mice triggers a systemic type-I IFN response and does not lead to a detectable activation of PKR or an UPR. Moreover, their chronic antiviral state protected *Adar*^Zα/Zα^ mice from virus infection.

### *Adar*^Zα/Zα^ mice develop a MAVS-dependent type-I IFN response

To test whether the spontaneous type-I IFN response caused by ADAR1 Zα-domain mutation depended on the MDA5/MAVS signalling pathway, we crossed *Adar*^Zα/Zα^ mice to animals that are deficient for MAVS. The mitochondrial outer membrane protein MAVS acts as a downstream signalling adaptor for the RIG-I like receptors (RLRs) RIG-I and MDA5 (Kawai et al., 2005; Meylan et al., 2005; Seth et al., 2005; Xu et al., 2005). As expected, 8-10 week old *Adar*^Zα/Zα^ mice displayed increased *Ifi44, Isg15* and *Zbp1* mRNA expression in lung, spleen and brain, however, ADAR1 Zα-domain mutant animals that were deficient for MAVS did not develop a spontaneous type-I IFN response (Figures 3A-3C). This provides genetic evidence that the enhanced type I IFN response due to loss Zα-domain mutation interaction with Z-RNA is driven by the RLRs RIG-I and/or MDA5. Based on our observation that mutation of the ADAR1 Zα-domain triggers an MDA5-mediated type-I IFN response in human cells (see Figure 1B) and the fact that the embryonic lethality of *Adar*^-/-^ mice is rescued in an MDA5-deficient background but not by RIG-I deficiency (Pestal et al., 2015), we conclude that the spontaneous antiviral immune response is mainly driven by the MDA5/MAVS signalling pathway.

**Figure 3.**
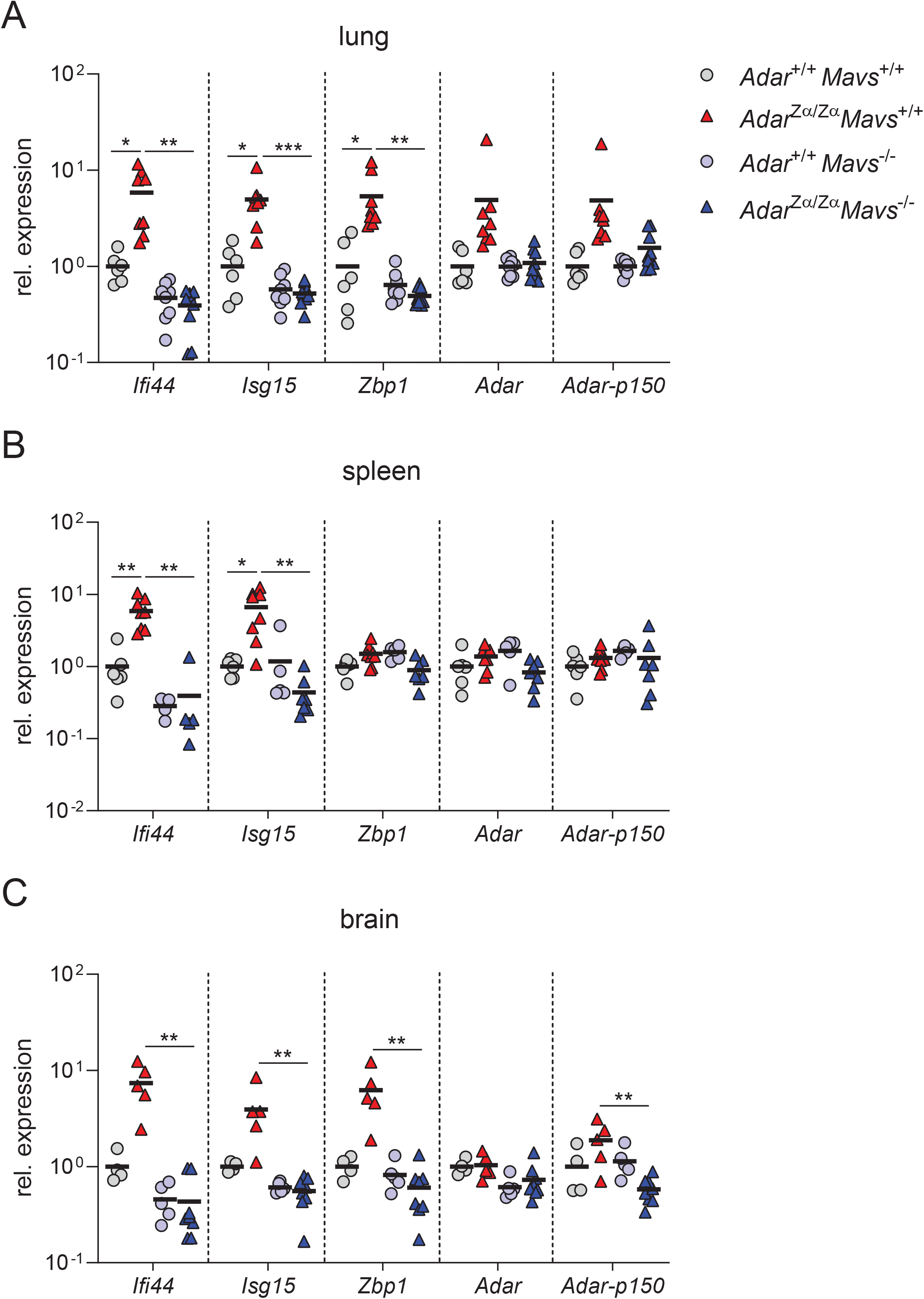
*Adar*^Zα/Zα^ mice develop a MAVS-dependent type-I IFN response. **(A-C)** RT-qPCR analysis of the indicated ISGs in lung (A), spleen (B) and brain (C) tissue from 8-10 week old wild type and ADAR1 Zα-domain mutant mice in a MAVS-sufficient (*Adar*^+/+^ *Mavs*^+/+^ or *Adar*^Zα/Zα^ *Mavs*^+/+^) and MAVS-deficient (*Adar*^+/+^ *Mavs*^-/-^ or *Adar*^Zα/Zα^ *Mavs*^-/-^) background. Data are pooled from two independent experiments. The mean relative expression of each gene in *Adar*^+/+^ *Mavs*^+/+^ tissues is set at 1. Lines represent the mean and symbols represent individual mice. *, P < 0.05. **, P <0.01, ***, P< 0.001 by unpaired t-test.

### *Adar*^Zα/Zα^ mice maintain normal haematopoiesis and do not develop autoinflammatory disease

Loss of ADAR1 activity results in severe defects in haematopoietic development, which is largely driven by spontaneous activation of MDA5 (Hartner et al., 2009; Liddicoat et al., 2016; Liddicoat et al., 2015). To determine whether haematopoiesis was affected in *Adar*^Zα/Zα^ mice, we performed whole blood phenotyping and detailed flow cytometry analysis of the leukocyte compartment of 8-10 week old animals. *Adar*^Zα/Zα^ mice were mildly anaemic as indicated by the reduced red blood cell numbers and haematocrit and haemoglobin levels compared to wild type littermates (Figure S4A). We did not detect any differences in the numbers of circulating lymphocytes (B cells, CD4 and CD8 T cells, and NK and NK T cells) or myeloid cells (neutrophils, basophils, eosinophils, and Ly-6C^-^ and Ly-6C^+^ monocytes) (Figure S4B and S4C). Together, this suggests that disrupted ADAR1/Z-RNA interaction has no effect on leukocyte cell development. Interestingly, Ly-6C^+^ monocytes expressed higher levels of CD64 also known as Fc-gamma receptor 1 (Figure 4A). The Interferome database categorises CD64 as an ISG and monocyte expression of CD64 strongly correlates with circulating type-I IFN levels in systemic lupus erythematosus (SLE) patients (Li et al., 2010). Moreover, anaemia is a common symptom of SLE (Aringer, 2020). Both the increased expression of CD64 and reduced presence of red blood cells therefore suggest that *Adar*^Zα/Zα^ mice may be prone to the development of SLE-like autoimmune disease.

**Figure 4.**
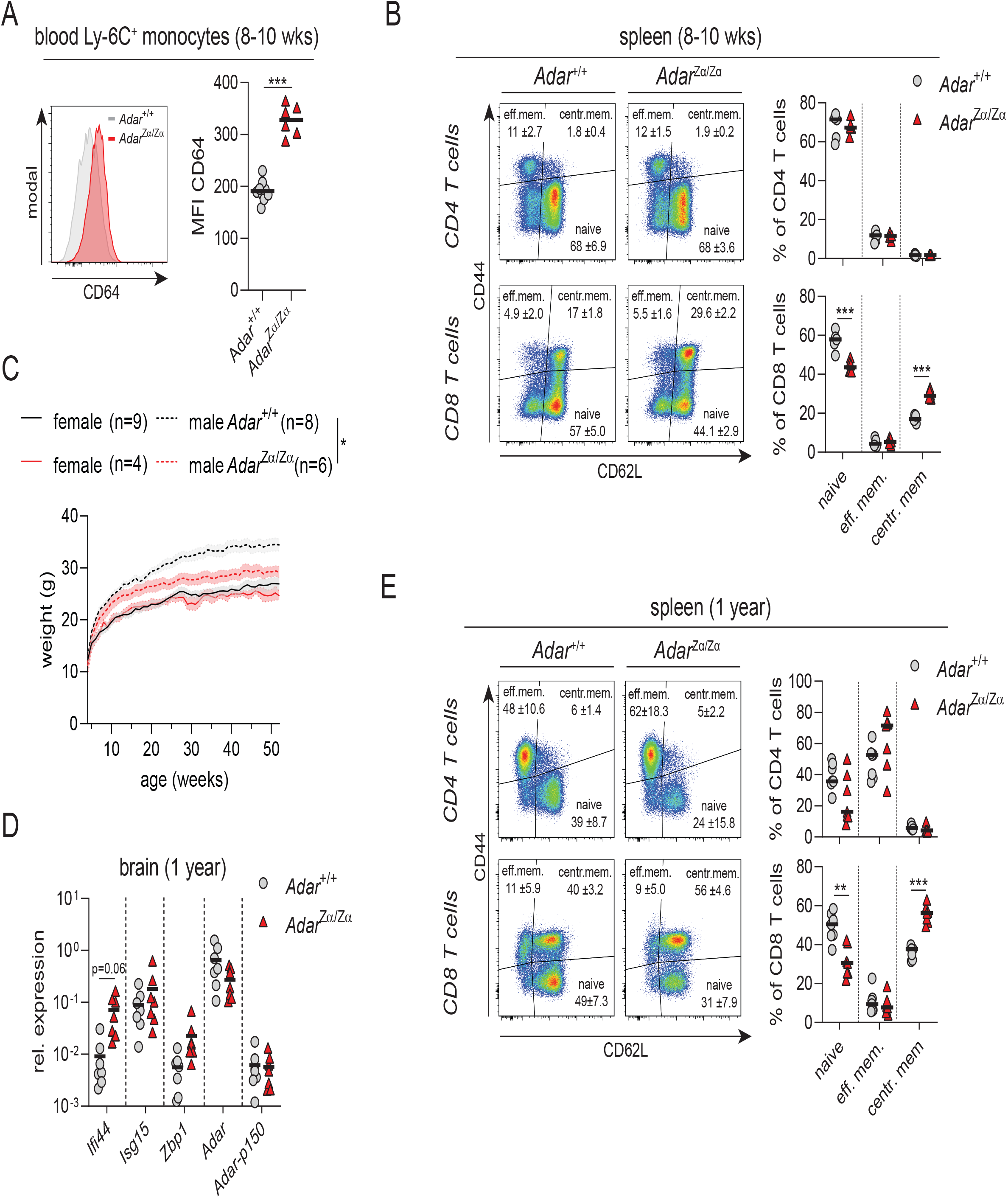
*Adar*^Zα/Zα^ mice maintain normal haematopoiesis and do not develop autoinflammatory disease. **(A)** Peripheral blood from 8-10 week old *Adar*^Zα/Zα^ mice and their wild-type littermates (*Adar*^+/+^) was analysed by flow cytometry. The histogram (left) depicts representative CD64 expression levels on Ly-6C^+^ monocytes. Quantification of CD64 expression levels on Ly-6C^+^ monocytes by median fluorescent intensity (MFI; right). The gating strategy is outlined in Figure S4B. Lines represent the mean and dots indicate individual mice. ***, P< 0.001 by Mann-Whitney U test). **(B)** Splenic CD4 and CD8 T cells from 8-10 week old mice from the indicated genotypes were analysed for CD44 and CD64L expression by flow cytometry. Gating of naive (CD44^-^CD62^lo^) and activated effector memory (eff. mem; CD44^hi^CD62^lo^) and central memory (centr. mem; CD44^+^CD62^hi^) CD4 and CD8 T cells are indicated by representative flow cytometry plots (left) for each genotype. Quantification of naïve and activated CD4 and CD8 T cell subpopulations as percentage of the total population (right). The gating strategy is outlined in Figure S4D. Lines represents the mean, and symbols depict individual mice. ***, P< 0.001 by unpaired t-test. **(C)** Weight in grams (g) of male and female mice of the indicated genotype was measured weekly from 4 until 52 weeks of age. Lines represent mean and shaded area shows ± SEM *, P < 0.05 by 2-way ANOVA. **(D)** RT-qPCR analysis of the indicated ISGs in brain tissue from the 1 year old *Adar*^+/+^ and *Adar*^Zα/Zα^ mice. Lines represents the mean and symbols depict individual mice. P = 0.06 by unpaired t-test. **(E)** Spleens from 1 year old mice of the indicate genotype were analysed as in (B). Lines represents the mean, and symbols depict individual mice. **, P < 0.01, ***, P < 0.001 by unpaired t-test. See also Figure S4.

Expansion of GL7^+^ CD95^+^ germinal center B cells and plasmablasts are indicative of antibody-mediated autoreactivity in a mouse model of SLE (Kool et al., 2011). Impaired function of Foxp3^+^ regulatory CD4 T cells (Tregs) and the accumulation of CD3ε^+^ CD4/CD8^neg^ B220^+^ nonconventional T cells are markers in several mouse models of autoimmune disease (Fontenot et al., 2003; Hori et al., 2003; Watanabe-Fukunaga et al., 1992). To test whether *Adar*^Zα/Zα^ mice developed SLE-like autoimmunity, we performed extensive flow cytometry-based phenotyping of the splenic T and B cell compartment (Figure S4D). We did not detect any notable increase in the number of GC B cells, plasmablasts, CD3ε^+^ CD4/CD8^neg^ B220^+^ T cells or a reduction in Tregs in the spleens of *Adar*^Zα/Zα^ mice. Despite a lack of these autoimmune markers, the spleens of ADAR1 Zα-domain mutant mice contained fewer naïve CD64L^hi^ CD44^-^ and more activated CD64L^hi^ CD44^+^ central memory CD8 T cells, while CD4 T cells were unaffected (Figure 4B).

Although 8-10 week old *Adar*^Zα/Zα^ mice did not display any signs of pathology, they showed inflammatory markers such as an elevated type-I IFN signature, anaemia and enhanced CD64 expression on monocytes accompanied by CD8 T cell activation. This suggests that they may develop autoimmune disease at later age. We therefore housed a cohort of *Adar*^Zα/Zα^ animals and littermate controls until 1 year of age. Despite the fact that *Adar*^Zα/Zα^ males were leaner than their wild type littermates (Figure 4C), we did not observe any macroscopic signs of disease. Instead, we found that the increase in ISG expression was less pronounced in the brains of 1 year old *Adar*^Zα/Zα^ mice compared to younger animals (Figure 4D; compare to figure 3C). In accordance with the loss of the type-I interferon signature in older animals we detected similar levels of CXCL10 protein in serum of 1 year old *Adar*^Zα/Zα^ and control littermates (Figure S4F). Aged ADAR1 Zα-domain mutant mice still showed enhanced CD8 T cell activation (Figure 4E), however, we did not observe an abnormal expansion of GC B cells, plasmablasts or CD3ε^+^ CD4/CD8^neg^ B220^+^ T cells or a reduction in Tregs in the spleens of these animals (Figure S4G).

Together these results show that *Adar*^Zα/Zα^ mice develop normally without a breach in T or B cell tolerance. The systemically elevated type-I IFN signature that is evident in 8 to 10 week old animals is not sustained until 1 year of age suggesting that a chronic exposure to type-I IFNs is well-tolerated until later in life without the development of autoinflammatory disease.

### Loss of ADAR1/Z-RNA interaction results in reduced A-to-I editing of SINEs

ADAR1 marks self-dsRNA by converting adenosine residues into inosines. Inosine-modified dsRNA is a poor agonist of MDA5 and the A-to-I editing activity of ADAR1 is essential to suppress an MDA5/MAVS-driven immune response (Ahmad et al., 2018; Liddicoat et al., 2015). To understand the molecular mechanism that drives the spontaneous type-I IFN response when the Z-RNA binding capacity of ADAR1 is abrogated, we asked whether A-to-I editing activity of ADAR1 is disrupted in Zα-domain mutant cells. A-to-I sites can be determined bioinformatically from RNA-Seq data as inosines are interpreted as guanosines by the reverse transcriptase during cDNA synthesis thereby introducing A-to-G (A>G) RNA-DNA-differences (RDDs) in the sequencing library. When aligned to the genomic sequence these A>G RDDs may specify a *bona fide* ADAR1 editing site.

We first determined A>G RDDs in the total stranded RNA-Seq dataset from mouse lung endothelial cells (see Figures 2F and 2G) isolated from *Adar*^Zα/Zα^ and control *Adar*^+/+^ mice using the machine-learning based algorithm RDDpred (Kim et al., 2016). We used previously annotated A-to-I sites from the REDIportal, DARNED and RADAR databases as positive training sets (Kiran et al., 2013; Mansi et al., 2020; Ramaswami and Li, 2014). We identified 20,534 unique A-to-I editing sites in mouse lung endothelium from the combined transcriptomes of 4 ADAR1 Zα-domain mutant and 4 wild type samples. Most of these sites were located in introns followed by 3’UTRs (Figure 5A). RepeatMasker analysis revealed that more than half of these sites mapped to repeat elements and that 60% of the repeat targets were located in short interspersed nuclear elements (SINEs), including B1, B2 and B4 repeats (Figure 5A). This is in agreement with previously determined A-to-I editing profiles of other mouse tissues (Licht et al., 2019; Neeman et al., 2006; Tan et al., 2017). From the 20,534 A-to-I sites identified in mouse lung endothelium, 13,590 targets were found in both Zα-domain mutant and wild type cells. We also identified 2,472 non-overlapping wild type-only sites and 4,472 sites that were only found in lung endothelium from *Adar*^Zα/Zα^ mice (Figure 5B). It is important to note that the likelihood of uncovering A-to-I sites is largely a function of sequencing depth (Bazak et al., 2014). This is especially problematic when dealing with sites that are edited at low levels and which would require a proportionally higher read coverage to enable their reliable detection. We can therefore not exclude that part of the sites identified as Zα-domain mutant-only or wild type-only were falsely assigned to these groups due to a lack in sequencing coverage. To avoid dealing with this bias we decided to only consider common sites that were detected in at least 2 wild type and 2 at least Zα-domain mutant samples. This resulted in the identification of 2,907 high-confidence A-to-I editing sites that occurred in both wild type and Zα-domain mutant samples (Figure 5B). To determine the impact of loss of ADAR1 binding to Z-RNA we measured the efficiency of ADAR1 editing by calculating the A-to-I editing index for each of these targets. The editing index is the number of reads containing an edited adenosine residue relative to the total number of reads covering that particular A-to-I site. Interestingly, we found that the overall editing efficiency was reduced in cells expressing the Zα-domain mutant variant of ADAR1 (Figure 5C). This was evident for sites covering the whole transcript including targets within 3’UTRs, introns and 5’UTRs and exons, annotated as “other” (Figure 5D). Further analysis showed that a larger fraction of sites had a lower editing index in most SINE family members, suggesting that loss of ADAR1/Z-RNA interaction led to an overall reduction in ADAR1 editing efficiency rather than causing a decrease in editing of a specific SINE subset (Figure S5B). Importantly, we did not observe any major differences in transcript abundance of different repeat sequence family members including the SINE/B1 and SINE/B2 elements between wild type and *Adar*^Zα/Zα^ genotypes, indicating that the reduced editing efficiency in Zα-domain mutant cells was not due to differential repeat expression (Figure S5A).

**Figure 5.**
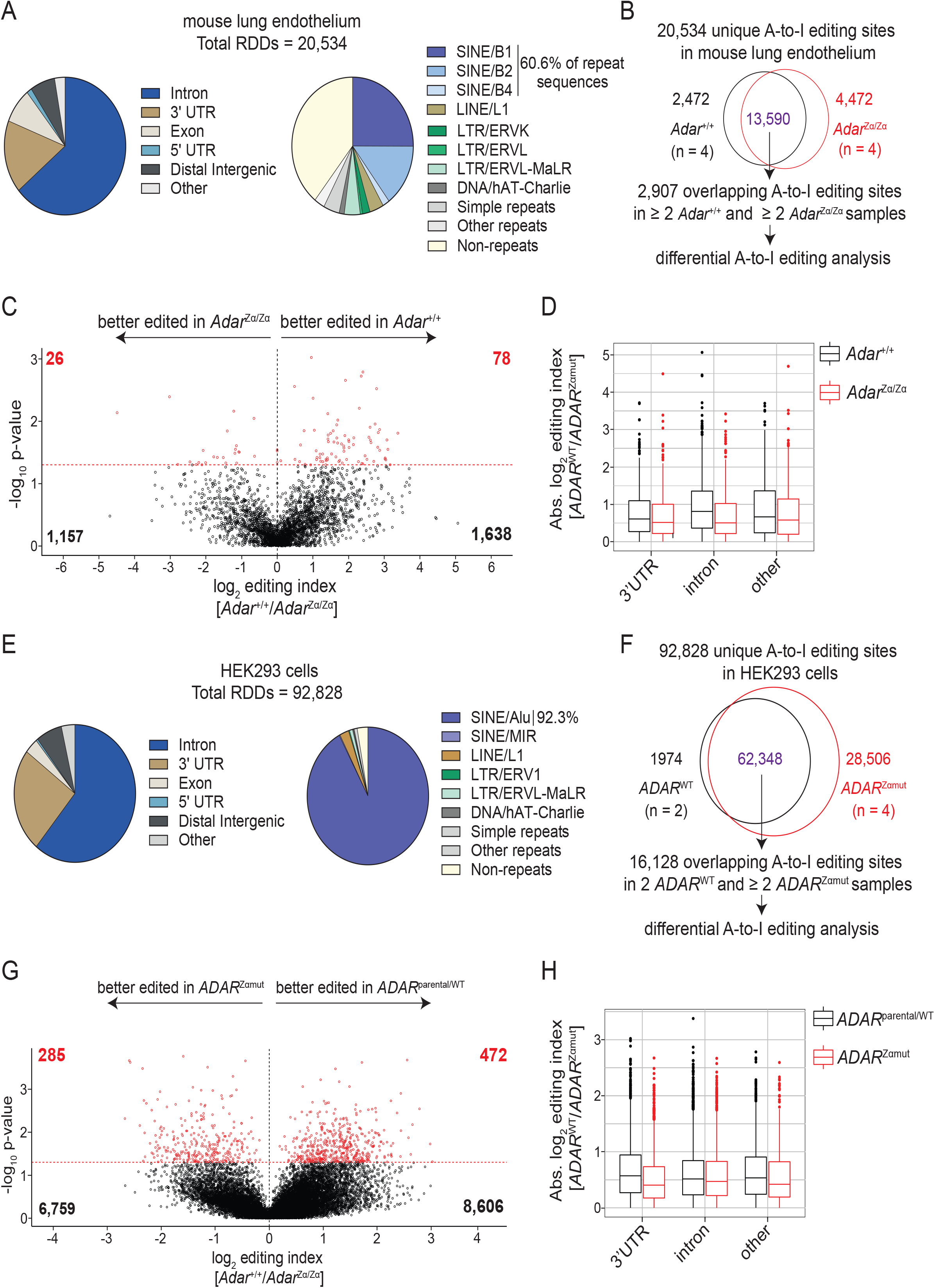
Loss of ADAR1/Z-RNA interaction results in reduced A-to-I editing of SINEs. **(A)** A total of 20,534 unique A-to-I editing sites (≥ 10 read coverage) were determined using total stranded RNA-Seq data obtained from FACS purified lung endothelial cells of *Adar*^+/+^ (n = 4) and *Adar*^Zα/Zα^ (n = 4) mice. Genomic (left) and RepeatMasker (right) annotation of the A-to-I sites are presented as pie charts. **(B)** Flowchart of the analysis pipeline to determine sites that are differentially edited between wild type and ADAR1 Zα-domain mutant transcriptomes. Differential editing analysis was performed on sites that were identified at least 2 *Adar*^+/+^ and at least 2 *Adar*^Zα/Zα^ samples. Volcano plot **(C)** and box plot **(D)** showing differential A-to-I editing in primary lung endothelial cells of *Adar*^+/+^ and *Adar*^Zα/Zα^ mice. The editing index of each site is the number of variant reads containing an A>G RDD by the total number of reads covering the site. Significance of differential editing [Y-axis in (C)] was determined by applying a Welch Two Sample t-test on the log10 editing index values of each sample for every site. The log2 editing index [X-axis in (C)] of each site was calculated by dividing the mean editing index in *Adar*^+/+^ samples by the mean editing index in *Adar*^Zα/Zα^ samples followed by log2 transformation. Positive values indicate a higher editing index in the wild type samples. The Y-axis in (D) displays the absolute editing index of sites that were better edited in wild type samples (black boxes) vs. Zα-domain mutant samples (red boxes) subdivided according to their genomic annotation. **(E)** A total of 92,828 unique A-to-I editing sites (≥ 10 read coverage) were determined using total stranded RNA-Seq data obtained from *ADAR*^WT/parental^ (n = 2) and *ADAR*^Zα/Zα^ (n = 4) HEK293 cells. Genomic (left) and RepeatMasker (right) annotation of the A-to-I sites are presented as pie charts. **(F)** Flowchart of the analysis pipeline to determine sites that are differentially edited between wild type and ADAR1 Zα-domain mutant transcriptomes. Differential editing analysis was performed on sites that were identified in *ADAR*^WT/parental^ and at least 2 *ADAR*^Zα/Zα^ samples. **(G)** Volcano plot and box plot **(H)** showing differential A-to-I editing in wild type and ADAR1 Zα-domain mutant HEK293 cells calculated using the same approach as in (C) and (D).

To align these observations to human cells, we performed total stranded RNA-Seq on 4 *ADAR*^Zαmut^, 2 *ADAR*^KO^, 1 *ADAR*^p150-KO^ HEK293 clone(s) and 2 control samples including a wild type clone and the parental HEK293 cells (see Figure S1B). The Zα-domain containing ADAR1-p150 isoform is lowly expressed in HEK293 cells and its expression is induced by type-I IFN treatment (see Figure 1D). To study the effect of Zα-domain mutation of ADAR1 on its A-to-I editing activity we therefore induced the expression of ADAR1-p150 by stimulating cells for 24 hours with IFN-α2. Using RDDpred we identified RDDs within each sample. The majority of the RDDs were A>G changes suggestive of an A-to-I editing site, but we also detected other permutations most notably C>T, G>A and T>C RDDs (Figure S5C). We refined our analysis by removing targets for which multiple RDDs were assigned and by subtracting false positive ADAR1 A-to-I editing sites occurring in the *ADAR*^KO^ clones. Finally, we removed RDDs that were annotated as single nucleotide polymorphisms in the published genomic sequence of the parental HEK293 cells (Lin et al., 2014) (Figure S5C). The resulting dataset largely contained high confidence A-to-I editing sites (A>G RDDs), while other RDDs were reduced to background levels. In line with previous findings (Chung et al., 2018; Pfaller et al., 2018), we found that overall A-to-I editing was reduced in *ADAR-p150*^KO^ cells, which do not express the Zα-domain containing ADAR1-p150 isoform, however, we identified an equal number of sites per million reads in *ADAR*^Zαmut^ and control cells (Figure S5C). We identified 92,828 unique A-to-I sites from the combined transcriptomes of 2 control and 4 *ADAR*^Zαmut^ samples. Similar to mouse endothelium, the majority of these sites mapped to introns and 3’UTRs and over 90% of editing sites were located in SINE/*Alu* repeats (Figure 5E), which is in agreement with other studies in human cells (Athanasiadis et al., 2004; Blow et al., 2004; Kim et al., 2004; Levanon et al., 2004). We identified 62,348 unique sites that were present in both wild type and ADAR1 Zα-domain mutant genotypes and 16,128 targets were present in the 2 control and at least 2 *ADAR*^Zαmut^ samples (Figure 5F). We then calculated the A-to-I editing index for each of these sites and found that – similar as in mouse cells – a larger fraction of targets was better edited in wild type cells than Zα-domain mutant HEK293 cells (Figure 5G). This was not a consequence of differential repeat expression between both genotypes, as ADAR1 Zα-domain mutation did not alter repeat RNA levels (Figure S5D). In contrast to our findings in mouse endothelium, we found that intronic sites were relatively unaffected by loss of Z-RNA interaction with ADAR1 while the reduction in editing efficiency was most pronounced in 3’UTRs and 5’UTRs and exons, annotated as “other” (Figure 5H). The reduction in overall editing could not be attributed to a loss of activity towards a single or multiple SINE/*Alu* family member(s) (Figure S5E).

Altogether, these results show that Z-RNA binding to ADAR1 promotes efficient A-to-I editing of SINEs. In human transcripts, introns remained relatively unaffected by loss of ADAR1 activity, while in mouse endothelium this affected the whole transcript. This suggests that the enhanced type-I IFN signature due to Zα-domain mutation of ADAR1 is driven by the recognition of SINE-derived duplex RNA by MDA5.

## Discussion

Around 1 million copies of the ∼300 bp SINE/*Alu* repeat element occupy the human genome thereby constituting around 11% of the total DNA content (Deininger, 2011; Lander et al., 2001). Human *Alu* elements and mouse B1 and B2 repeats are enriched in genes and are especially prevalent in introns and 3’UTRs (Lu et al., 2020). Given their sheer abundance, SINEs are thought to have a profound impact on cell function and direct processes such as transcription, epigenetic control and gene evolution (Burns and Boeke, 2012; Lee et al., 2015). It is estimated that one new *Alu* insertion occurs every 20 births (Cordaux et al., 2006). The acquisition of SINEs inside genes may generate genomic plasticity, however, it comes at the risk of invoking an unwanted immune response (Kassiotis and Stoye, 2016; Mu et al., 2016). Transcripts that contain two adjacent and oppositely oriented SINEs can form dsRNA structures. Computational analysis on the orientation of repeat elements and their likelihood to form dsRNA across species, revealed that there is a strong selection against the presence of neighbouring inversely oriented repeat sequences in the 3’UTR of mRNAs, but not in intron-containing pre-mRNAs (Barak et al., 2020). Purifying selection against dsRNA structures is incomplete and inverted SINE repeats that have escaped this process may activate MDA5. Ultradeep sequencing revealed that nearly every adenosine in *Alu* elements with potential to form dsRNA is subject to the A-to-I editing activity of ADAR1 (Bazak et al., 2014). ADAR1 activity thus forms a second barrier to prevent deleterious MDA5 activation by SINE-derived dsRNA molecules.

The importance of ADAR1 in supressing MDA5-mediated immunity is evident from AGS patients with loss-of-function mutations in *ADAR*, which develop a pathogenic type-I IFN response (Rice et al., 2012). In this study, we focused on the observation that over half of *ADAR* AGS patients carried the p.Pro193Ala mutation in combination with a dysfunctional *ADAR* variant (Rice et al., 2017). Proline 193 lies within the Z-RNA binding Zα-domain of the p150-isoform of ADAR1 and its mutation is predicted to disrupt ADAR1/Z-RNA interaction. These patients thus express a mutant ADAR1 protein that is unable to bind to Z-RNA, but retains the capacity to bind to A-form dsRNA and has fully functional A-to-I editase domain. This implies that the interaction between the poorly defined Z-conformer of dsRNA and ADAR1 is required to fully exert its immunosuppressive effects. Indeed, Zα-domain mutation led to reduced ADAR1 activity measured by an editing reporter plasmid and *in vitro* ADAR1-p150 activity was enhanced towards a dsRNA substrate containing an alternating CG-sequence that is prone to form Z-RNA (Koeris et al., 2005; Wong et al., 2003). We here performed a transcriptome-wide analysis of thousands of A-to-I sites in human and mouse cells that express Zα-domain mutant ADAR1 and found that overall editing efficiency was reduced by loss of ADAR1/Z-RNA interaction. While the reduction in ADAR1 activity in mouse cells applied to whole transcripts, this was particularly evident for 3’UTRs in human transcripts. As introns are spliced out before nuclear export, SINE-derived dsRNA structures in 3’UTRs most likely form the immunostimulatory endogenous agonist of MDA5, which localises to the cytosol. The Zα-domain containing ADAR1-p150 isoform shuttles between the nucleus and the cytosol, hence an important part of its immunosuppressive may be to modify dsRNA structures in 3’UTRs of mRNAs that were recently exported out of the nucleus.

The identity of the RNA sequences that form Z-RNA remains unknown. The capacity of dsRNA to transition into the thermodynamically unfavourable Z-conformation is dependent on the sequence context. A 6 base pair r(GC)3 RNA duplex forms the minimum Zα-domain interaction motif, at least *in vitro* (Placido et al., 2007). The Zα-domain of ADAR1 could in theory bind to any dsRNA structure containing such a motif. Indeed, putative Z-RNA forming sequences were annotated in *Alu* repeats (Herbert, 2019). Moreover, recent studies in mice reported that SINE-derived duplex RNA may serve as an endogenous ligand for the Z-RNA sensor ZBP1, the only mammalian protein aside from ADAR1 that contains a Zα-domain (Jiao et al., 2020; Wang et al., 2020). We were not able to discern one or more members of the human *Alu* and mouse B1/B2 elements that were particularly “hypo-edited” due to Zα-domain mutation and that could point to a particular SINE-sequence containing a Zα-domain binding motif. To determine these Z-RNA-forming motifs, we would need to identify each SINE in relation to its inversely oriented pairing partner. It is possible that a particular combination of paired SINEs family members is more prone to form a Z-RNA structure. To identify these sequences one could employ the recently developed MDA5 RNase protection/RNA-Seq method (Ahmad et al., 2018) or perform ADAR1 CLIP-Seq in ADAR1 Zα-domain mutant cells.

Homozygous *Adar*^Zα/Zα^ mice display a subclinical type-I IFN response and do not develop AGS-like pathology even after 1 year of age. AGS patients express the Pro193Ala ADAR1 variant only from one allele, suggesting that gene dosage may determine whether pathology develops. Crossing *Adar*^Zα/Zα^ mice to ADAR1-deficient mice to generate *Adar*^Zα/null^ that express half the amount of ADAR1 Zα-domain mutant protein compared to homozygous *Adar*^Zα/Zα^ animals may be an interesting approach to test this hypothesis. The missense *ADAR* p.Pro193ala mutation has an unusually high allele frequency of ∼0.2%, suggesting that this variant may offer an evolutionary advantage. We found that *Adar*^Zα/Zα^ mice were more resistant to virus infection. It is possible that heterozygous carriers of the p.Pro193ala allele produce slightly elevated levels of type-I IFNs to better deal with viral insults. Genetic lesions resulting in subclinically elevated type-I IFN levels may thus supplement the microbiota-driven tonic type-I IFN response and prime the immune system for future infections (Abt et al., 2012; Bradley et al., 2019; Ganal et al., 2012; Schaupp et al., 2020; Steed et al., 2017; Winkler et al., 2020).

In sum, we provide genetic evidence that Z-RNA binding to ADAR1 is crucial to maintain tolerance to self-RNA. We propose that ADAR1/Z-RNA interactions widens the editing repertoire of ADAR1 and that disruption of this interaction leads to the accumulation of immunostimulatory SINE-derived dsRNA that may trigger the onset of human autoinflammatory diseases such as AGS.

## Acknowledgements

We would like to thank Wim Declercq for helpful discussions and critical reading of the manuscript. We are grateful to A. Wullaert for providing *Mavs*^-/-^ mice, to E. Picardi for providing the mouse A-to-I editing site positive training dataset for RDDpred (Licht et al., 2019), the VIB Protein Core for recombinant Cas9-GFP and to the VIB-UGent IRC Animal house for mouse husbandry. We would like to thank the VIB Flow Core for training, support and access to the instrument park. This research would not have been possible without support from the following funding agencies. R. de Reuver was supported by a Ghent University BOF PhD fellowship (01D24020). L. Seys was supported by an FWO postdoctoral fellowship (1236420N). Research in the Vandenabeele group is supported by EOS MODEL-IDI (FWO Grant 30826052), FWO research grants (G.0E04.16N, G.0C76.18N, G.0B71.18N, G.0B96.20N), Methusalem (BOF16/MET_V/007), Foundation against Cancer (F/2016/865, F/2020/1505), CRIG and GIGG consortia, and VIB. J. Maelfait was supported by and Odysseus II Grant (G0H8618N) from the Research Foundation Flanders and by Ghent University.

## Author contributions

Conceptualisation, R.d.R., E.D. and J.M. Methodology, R.d.R., E.D., B.W., K.S. and F.V.N., J.M. Software, R.d.R., T.M. and A.B. Formal analysis, R.d.R. Investigation, R.d.R., E.D., B.W., K.S., L.S., E.D.M. and J.M. Resources, B.L., F.V.N., P.V. and J.M. Writing – Original Draft, R.d.R. and J.M. Writing – Review & Editing, R.d.R. and J.M. Funding Acquisition, R.d.R. and J.M. Supervision, J.M.

## Materials and Methods

### Mice

*Adar*^fl-Zα^ mice were generated in C57Bl/6 ES cells by Cyagen and were kindly provided by Jan Rehwinkel. An *Adar*^+/fl-Zα^ male was crossed to a female Sox2-Cre deleter mouse to generate *Adar*^+/Zα^ offspring. Adar^+/Zα^ mice were interbred to generate homozygous ADAR1 Zα-domain mutant (*Adar*^Zα/Zα^) animals. *Adar*^+/+^ littermates were used as controls. Sox2-Cre mice were generated in CBA ES cells (Hayashi et al., 2003) and were purchased from The Jackson Laboratory (B6.Cg-*Edil3*^*Tg(Sox2-cre)1Amc*^/J). Sox2-Cre animals were backcrossed at least 10 times to a C57Bl/6 background. *Mavs*^-/-^ mice were generated in C57Bl/6 ES cells and were obtained from the Jurg Tschopp laboratory (Michallet et al., 2008). Mice were housed in individually ventilated cages at the VIB-Ugent Center for Inflammation Research in a specific pathogen-free facility, according to national and institutional guidelines for animal care. All experiments were conducted following approval by the local Ethics Committee of Ghent University.

### Cell culture

HEK293A, hereafter referred to as HEK293, cells (Lin et al., 2014) were cultured at 37°C and 5% CO2 in DMEM/F-12 (Dulbecco’s Modified Eagle Medium/Nutrient Mixture F-12, Gibco;31330-038) supplemented with 10% FCS (Bodinco), 2mM glutamine (Lonza; BE17-605F) and 1mM sodium pyruvate (Sigma-Aldrich; S8636). For bone marrow-derived macrophages (BMDMs), femurs and tibia were collected of 8-10-week-old *Adar*^Zα/Zα^ mice and wild type littermates. Bones were sterilised in 70% ethanol before flushing them with RPMI 1640 medium using a 26G needle. Next, cells were homogenized and filtered through a 70 µm cell strainer, followed by erythrocyte lysis with ACK Lysing Buffer (Lonza; 10-548E). Bone marrow cells were plated in culture medium containing 20% conditioned L929 supernatant and medium was refreshed after 3 days and cells were used for experiments 6 days after isolation. BMDMs were cultured at 37°C and 5% CO2 in RPMI 1640 medium supplemented with 10% FCS, 2mM glutamine (Lonza; BE17-605F), 1mM sodium pyruvate (Sigma-Aldrich; S8636), 100 U/ml penicillin, 100 µg/ml streptomycin (Sigma-Aldrich; P4333) and 20% conditioned L929 supernatant. Primary lung fibroblasts were isolated from lungs of 8-10-week-old male *Adar*^Zα/Zα^ mice and wild type littermates. Lungs were sterilized with 70% ethanol before cutting the tissue in pieces of ∼25mm. Digestion was performed with a 0.1% collagenase D (Roche;11088866001) and 0.2% trypsin solution (Gibco;15090-046) for 30 min at 37°C. The collagenase D / trypsin solution was refreshed and digestion continued another 30 minutes at 37°C. After neutralization with αMEM-20 [αMEM, Gibco; 22571-020, supplemented with 20% FCS, 2mM glutamine (Lonza; BE17-605F), 1mM sodium pyruvate (Sigma-Aldrich; S8636), 100 U/ml penicillin and 100 µg/ml streptomycin (Sigma-Aldrich; P4333)], cells were pelleted at 400g for 5 min and resuspended in αMEM-20 culture medium. The purity of primary lung fibroblasts was assured by passaging each fibroblast line for at least 3 times in αMEM-20, which selectively supports growth of fibroblasts, whereas other cell types die or stop proliferating. Primary murine lung fibroblasts were maintained under hypoxic conditions (3% O2 at 37°C and 5% CO2 in αMEM-20).

### Generation of *ADAR*^Zαmut^, ADAR^p150-KO^ and *ADAR*^KO^ HEK293 clones

Cells were electroporated (NEPA21, NepaGene) with the indicated gRNAs and GFP-tagged Cas9 (VIB Protein Service Facility). *ADAR*^KO^ clones were generated using a gRNA#1 targeting against exon 2, which is shared by the ADAR1 p110 and p150 isoforms. The *ADAR*^p150-KO^ clone was generated using gRNA#2 and gRNA#3 directed against exon 1A containing the p150-specific transcription start site. *ADAR*^Zαmut^ domain mutant cells were generated by CRISPR/Cas9-mediated homology directed repair and using gRNA#4. A ssDNA repair template oligo containing the N173A/Y177A mutations was electroporated with the Cas9/gRNA complex. 24 hours after electroporation, single GFP-positive cells were FACS purified and plated into 96-well plates. Clones were screened by PCR and protein expression was analysed by Western blot. Zα-domain mutation in *ADAR*^Zαmut^ clones was confirmed by Sanger sequencing and subcloning of the PCR fragments used for sequencing with the Zero Blunt® TOPO® PCR Cloning Kit (Invitrogen; 450245). At least 11 subclones were individually sequenced (see Figure S1A). Sequences of the gRNAs, ssDNA repair template and PCR primers are provided in the Key Resource Table.

### Reagents

Mouse CXLC10 protein was quantified using ELISA (eBioscience; BMS6018MST) and Bio-Plex (Bio-Rad; #12002244). Human recombinant IFN-α2 was from Biolegend (592704) and hybrid IFNα-B/D was from Novartis (CGP35269).

### Luciferase assays

HEK293 cells were seeded at a density of 200,000 cells per well of a 24-well plate and transfected 16h later with 200 ng of the specified expression vectors (see Key Resources Table) using Lipofectamine 2000 (Invitrogen; 11668-027). For titration experiments, empty vector was added to the transfection mix to bring the total of input DNA to 200 ng. 50 ng of an IFNβ promoter reporter plasmid (p125-luc) and 20 ng of a *Renilla* luciferase reporter (pRL-TK; Promega E2241) under control of a thymidine kinase promoter were co-transfected. 24h after transfection, cells were washed with PBS and lysed in Passive Lysis Buffer, 5X (Promega; E194A). Luciferase units were measured by the Dual luciferase reporter assay system (Promega; E1910) on a Promega GloMax luminometer. Relative luciferase units (RLU) were calculated by dividing normalised luciferase units (Firefly / *Renilla*) by the normalised luciferase units from cells transfected with 200 ng empty vector for each HEK293 clone.

### Western Blotting and Cellular Fractioning

Fractioning of primary murine lung fibroblasts was performed with the Cell Fractionation Kit of Cell Signaling Technology (#9038S). For Western blotting, cells were washed with ice-cold PBS and lysed in protein lysis buffer (50mM Tris HCl, pH 7.5, 1% Igepal CA-630 and 150mM NaCl) supplemented with complete protease inhibitor cocktail (Roche; 11697498001) and PhosSTOP (Roche;4906845001). Lysates were cleared by centrifugation at 12,000 g for 15 min and 5x Laemlli loading buffer (250mM Tris HCl, pH 6.8, 10% SDS, 0.5% Bromophenol Blue, 50% glycerol and 20% β-mercaptoethanol) was added to the supernatant. Finally, samples were incubated at 95°C for 5 min and analysed using Tris-Glycine SDS-PAGE and semi-dry immunoblotting. Primary and secondary antibodies for Western blotting are listed in the Key Resources Table.

### Quantitative RT-PCR (RT-qPCR)

Tissue sections were homogenized with the Precellys 24 Tissue Homogeniser and 1.4 mm Zirconium oxide beads (VWR International; P000927-LYSK0-A) in Trizol reagent (Gibco BRL; 15596-018). For both tissues and cells, total RNA was purified using RNeasy colums (Qiagen) with on-column DNase I digestion. cDNA synthesis was performed with the SensiFast cDNA synthesis kit (Bioline; BIO-65054). SensiFast SYBR No-Rox kit (Bioline; BIO-98005) or PrimeTime qPCR Master Mix (IDT; 1055771) was used for cDNA amplification using a Lightcycler 480 system (Roche). Primers and probes are listed in the Key Resources Table.

### Tissue and peripheral blood processing for flow cytometry

Lungs and spleens of 8-10 week old *Adar*^Zα/Zα^ and wild type littermates were collected in ice-cold PBS. Lungs were processed according to (Fletcher et al., 2011) with minor modifications. In brief, lungs were digested for 1 hour at 37°C in digestion buffer [RPMI 1640 supplemented with 10% FCS, 0.8 mg/ml dispase II (Invitrogen; 17105-041), 0.2 mg/ml collagenase P (Sigma-Aldrich; 000000011213857001) and 0.1% DNase I (Roche; 1010459001)] with regular mixing. After neutralisation with FACS buffer without sodium azide (PBS, 5% FCS, 1 mM EDTA), erythrocytes were lysed with ACK Lysing Buffer (Lonza; 10-548E) and filtered through a 100 µm cell strainer to obtain single cell solutions. Next, cells were stained with anti-mouse CD16/CD32 (Fc-block; BD Biosciences; 553142) and anti-mouse CD45-FITC before magnetic separation using anti-FITC microbeads (Miltenyi Biotec; 130-048-701) on midiMacs type LS separation columns (Miltenyi Biotec; 130-042-401). Spleens were digested for 30 min at 37°C in digestion buffer [(RPMI 1640 supplemented 0.5 mg/ml collagenase D (Roche; 11088866001) and 10 µg/ml DNase I (Roche; 1010459001)] with regular mixing. After neutralisation with RPMI 1640 medium containing 2% FCS, erythrocytes were lysed with ACK Lysing Buffer (Lonza; 10-548E) and filtered through a 70 µm cell strainer to obtain single cell solutions. Peripheral blood was collected in EDTA-coated tubes (Sarstedt; 20.1288) by tail vein bleeding. 1 ml ACK Lysing Buffer (Lonza; 10-548E) was mixed with 100 µl blood and incubated for 10 min at RT. Cells were washed 2 times with ice-cold PBS and were stained for flow cytometry analysis.

### Flow cytometry

Single cell suspensions were first stained with anti-mouse CD16/CD32 (Fc-block; BD Biosciences; 553142) and dead cells were excluded with the Fixable Viability Dye eFluor506 (eBioscience; 65-0866-14) for 30 min at 4°C in PBS. Next, cell surface markers were stained for 30 min at 4°C in FACS buffer (PBS, 5% FCS, 1mM EDTA and 0.05% sodium azide). Cells were acquired on an LSR Fortessa or a FACSymphony (BD Biosciences) or sorted on a Aria III sorter (BD Biosciences). Data were analysed with FlowJo software (Tree Star). The total number of cells was counted on a FACSVerse (BD Biosciences). Fluorochrome-conjugated antibodies are listed in the Key Resources Table.

### Blood analysis

Peripheral blood was collected in EDTA-coated tubes by tail vein bleeding. Total number of leukocytes, erythrocytes whole-blood parameters were determined using a HEMAVET^®^950LV (Drew Scientific).

### EMCV infections

8-10 week old *Adar*^Zα/Zα^ and wild type littermates were infected by intraperitoneal injection with 200 or 2000 plaque forming units (pfu) of Encephalomyocarditis virus (ATCC® VR-129B(tm)). 48 or 72 hour post-infection, hearts were homogenized with a Precellys 24 Tissue Homogeniser and 1.4 mm Zirconium oxide beads (VWR International; P000927-LYSK0-A) in 1 mL of Trizol reagent (Gibco BRL; 15596-018), followed by total RNA extraction as described above.

### RNA-Seq library preparation

Total RNA of IFN-α2 treated HEK293 cells was purified using RNeasy columns (Qiagen) with on-column DNase I digestion. RNA integrity was tested with the Agilent RNA 6000 Pico Kit (Agilent; 50671513). Sequencing libraries were prepared using the Truseq Stranded Total RNA kit (Illumina) with the Ribo-Zero Gold kit (Illumina) for ribosomal depletion. Total RNA of primary murine lung endothelial cells was purified using RNeasy Micro columns (Qiagen) with on-column DNase I digestion. RNA integrity was checked with the Agilent RNA 6000 Pico Kit (Agilent; 50671513). Libraries were prepared using the SMARTseq Stranded Total RNA kit (Takara Bio) with ZapR kit (Takara Bio) for ribosomal depletion. Libraries were sequenced on an Illumina HiSeq3000 genreating 150 bp paired-end sequencing reads.

### Differential Gene Expression Analysis

Quality control of raw Fastq files was performed with FastQC v0.11.7. Trimmomatic v1.4 was used for clipping of adapter and index sequences with default parameters. Paired-end reads were mapped to the murine reference genome (GRCm38) using STAR 2.7.3a in 2-pass mode. Raw read counts were determined using HTSeq 0.11.1 in union mode. Normalised counts were calculated with R-package DESeq2 (v1.26.0). Gene ontology enrichment was analysed using DAVID (https://david.ncifcrf.gov/); functional annotation; ENSEMBLE_GENE_ID; GO-term BP direct; Bonferonni EASE = 0.05.

### A-to-I editing analysis

In addition to adapter and index sequence clipping, Trimmomatic v1.4 was used to crop the first 15 bp and last 6 bp of the reads to circumvent bias introduced by random priming. Trimmed paired-end reads were mapped to the reference genome (HEK293; assembly hg19, murine lung endothelial cells; assembly mm10) using STAR 2.7.3a. Obtained BAM files (sorted by coordinate), the reference genome and machine learning training sets functioned as input files for the RDDpred tool (v1.1) (Kim et al., 2016). RDDpred was used for the detection of RNA-DNA differences. Training sets consisted of Mapping Error-prone Sites (negative set; http://epigenomics.snu.ac.kr/RDDpred/prior.php). and annotated A-to-I editing sites (positive set) derived from the DARNED (Kiran et al., 2013), RADAR (Ramaswami and Li, 2014) and REDIportal (Mansi et al., 2020) databases. Murine editing sites derived from the RADAR database were converted from mm9 to mm10 assembly using LiftOver (Kuhn et al., 2013). To assign strand topology to each identified RNA-DNA differences (RDDs), BAM files were split in half-samples containing reads that were either mapped to the sense or antisense strand. Bam-readcount (https://github.com/genome/bam-readcount) was used to determine read coverage and variant frequency for all RDDs in each half-sample. Ambiguous RDDs (e.g. multiple variant calls per site, RDD detected on both strands), were excluded from downstream analysis. In addition, sites overlapping with annotated C57Bl/6-specific SNPs extracted from the Sanger Institute Mouse Genomes project v3 (dbSNPv137) and sites overlapping annotated SNPs were removed from further downstream analysis. HEK293-specific SNPs were retrieved from http://hek293genome.org/v2/data.php; data track CG 293 and converted to hg19 assembly using LiftOver. For HEK293 samples, an additional filtering step based on RDDs that were also detected in the ADAR1 KO sample was performed. Genomic annotation of the identified A-to-I editing sites was assigned using GENCODE basic annotation (HEK293; release 34, mouse: release 25, hierarchy: 3’ UTR > 5’ UTR > Exon > Intron > Intergenic > Downstream > Promoter) and repeat element status was determined based on RepeatMasker annotation (www.repeatmasker.org; Repeat Library 20140131). For differential editing analysis, the editing index was calculated (variant count / total count) and sites with a read count below 10 or editing index of 0 were discarded. Differentially edited sites (p-value <0.05) were determined by the Welch Two Sample t-test log10 values of the calculated editing index.

### Statistical analyses

Statistical analyses were performed using Prism 8.2.1 (GraphPad Software). Statistical methods are described in the figure legends.

## Key Resources Table

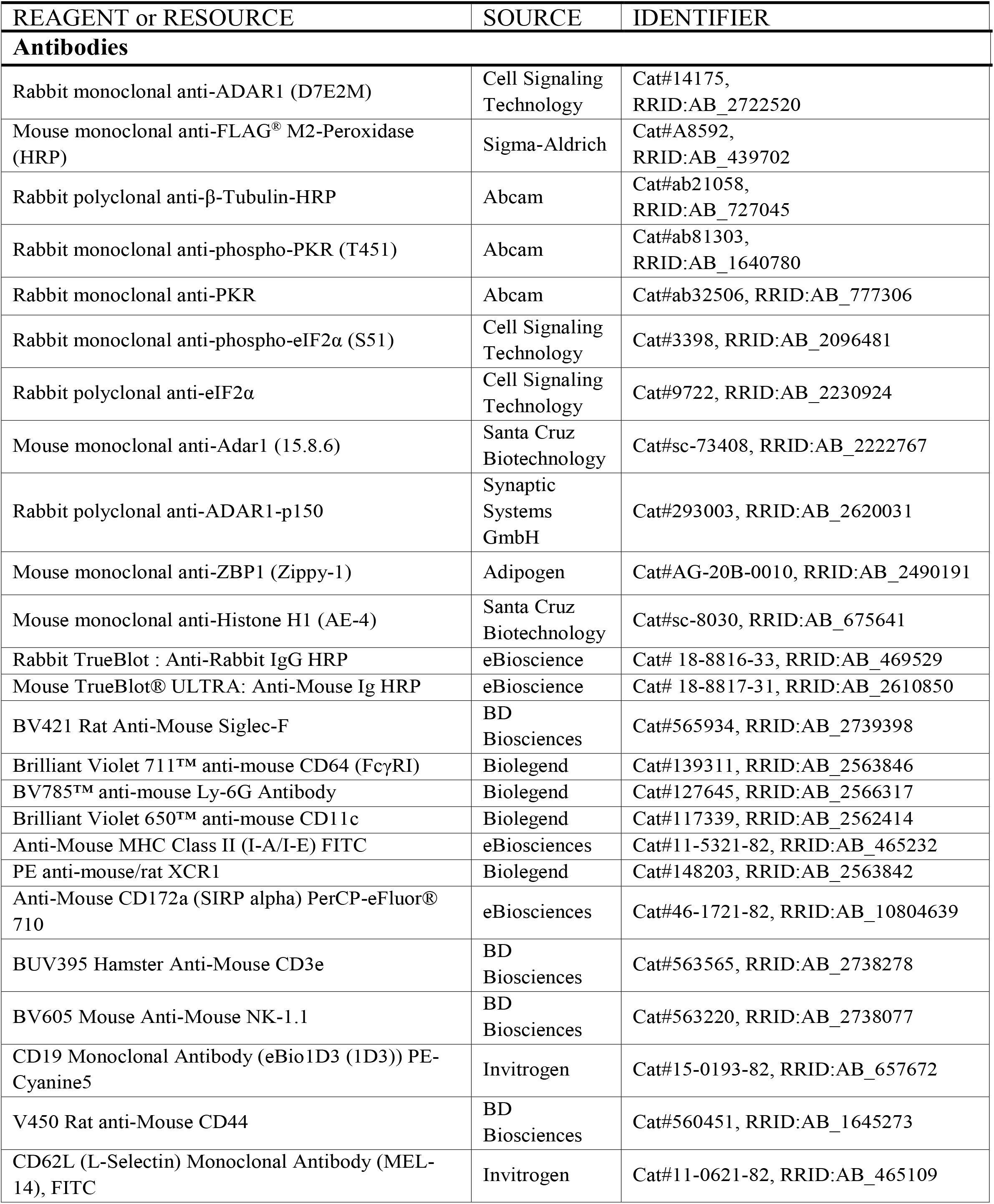
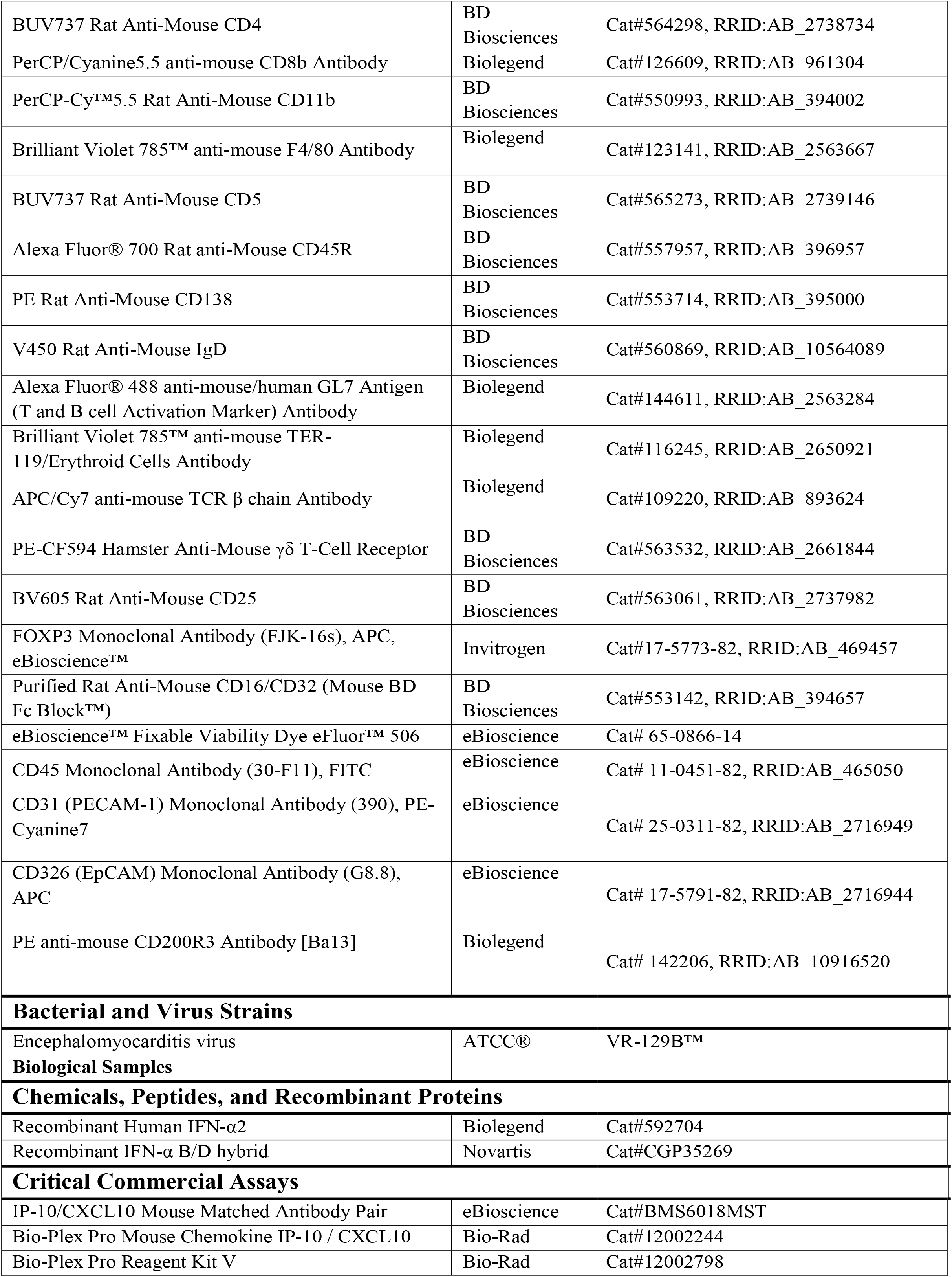
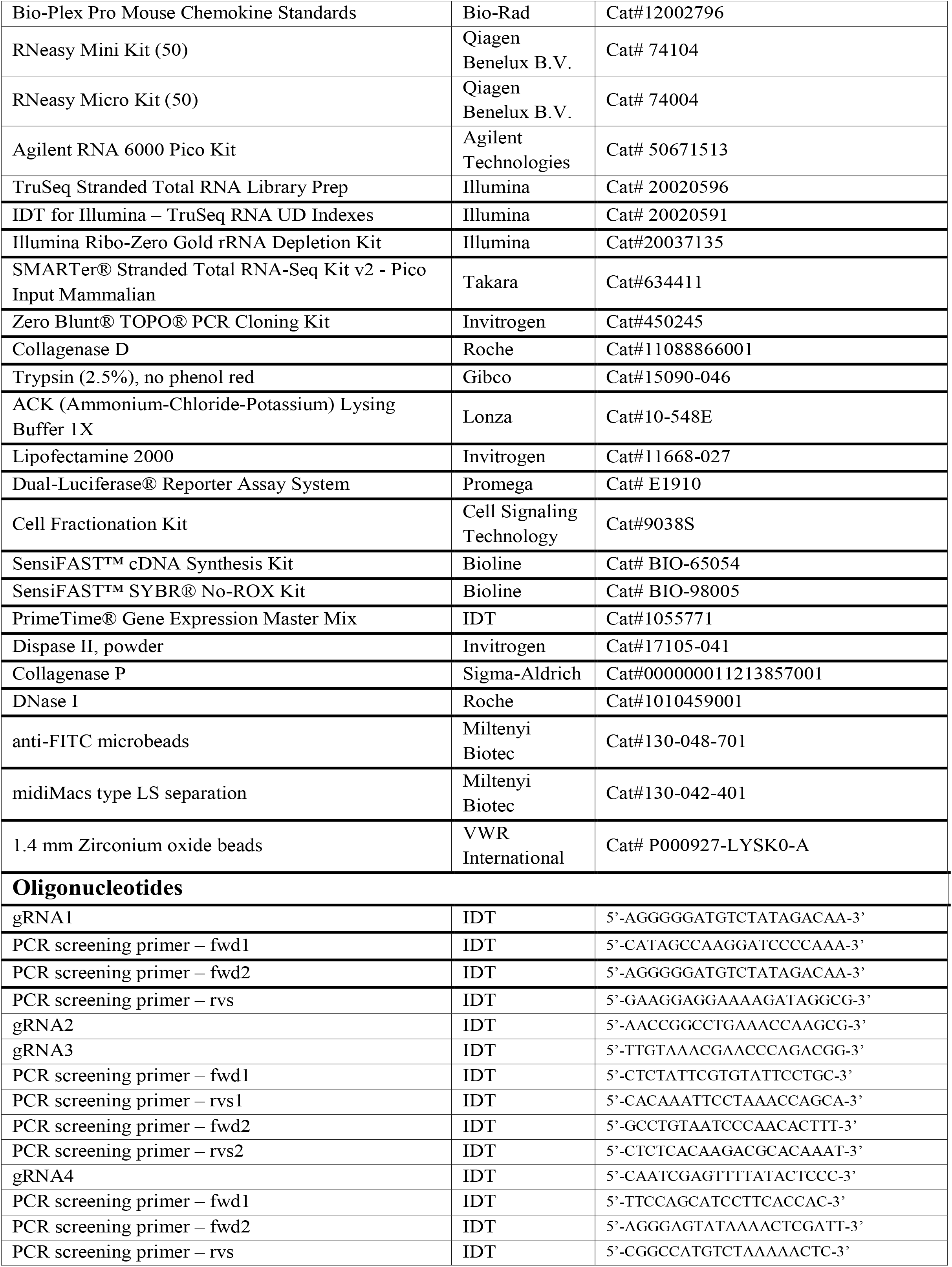
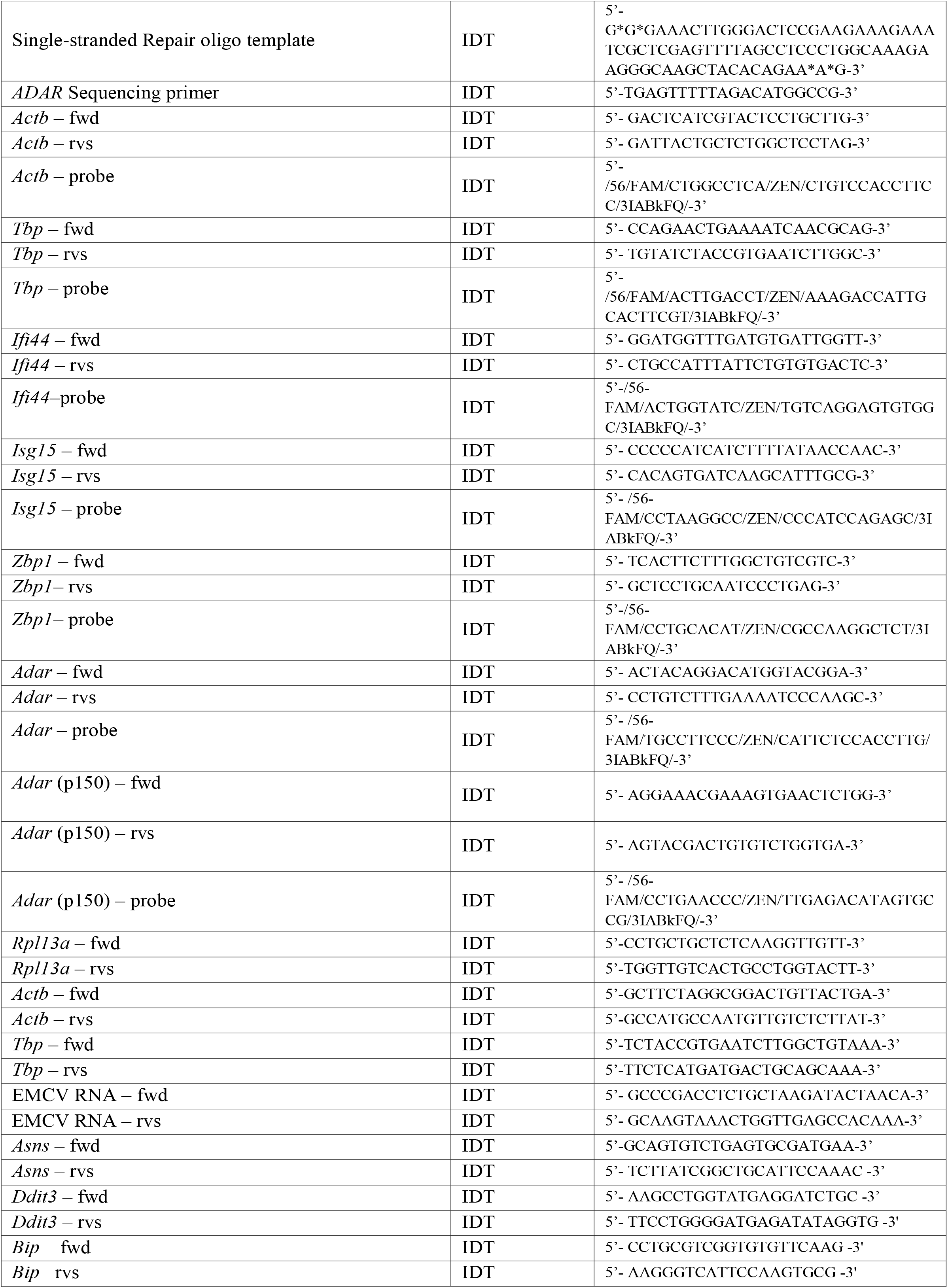
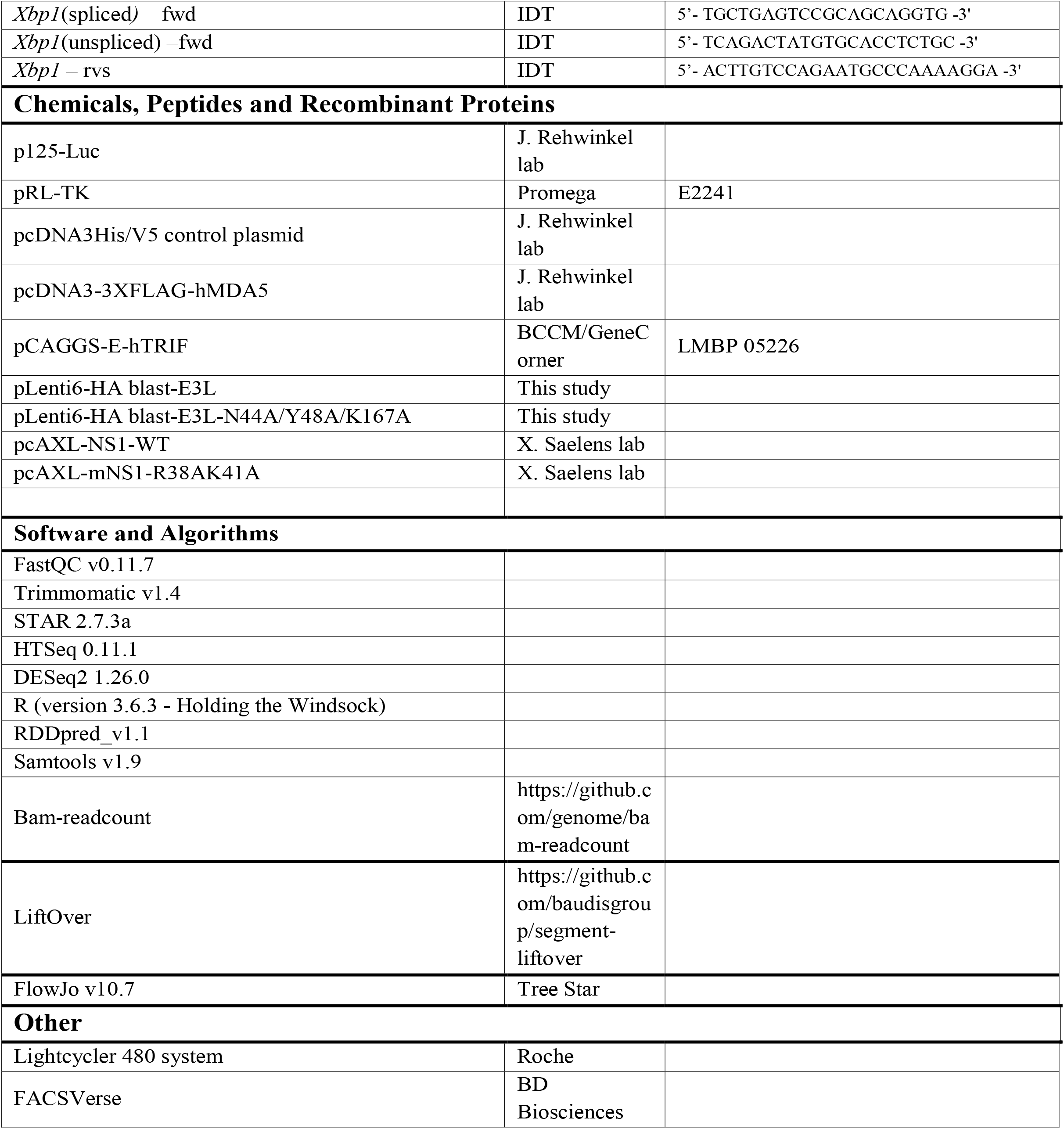

---

**Supplemental Figure 1.**
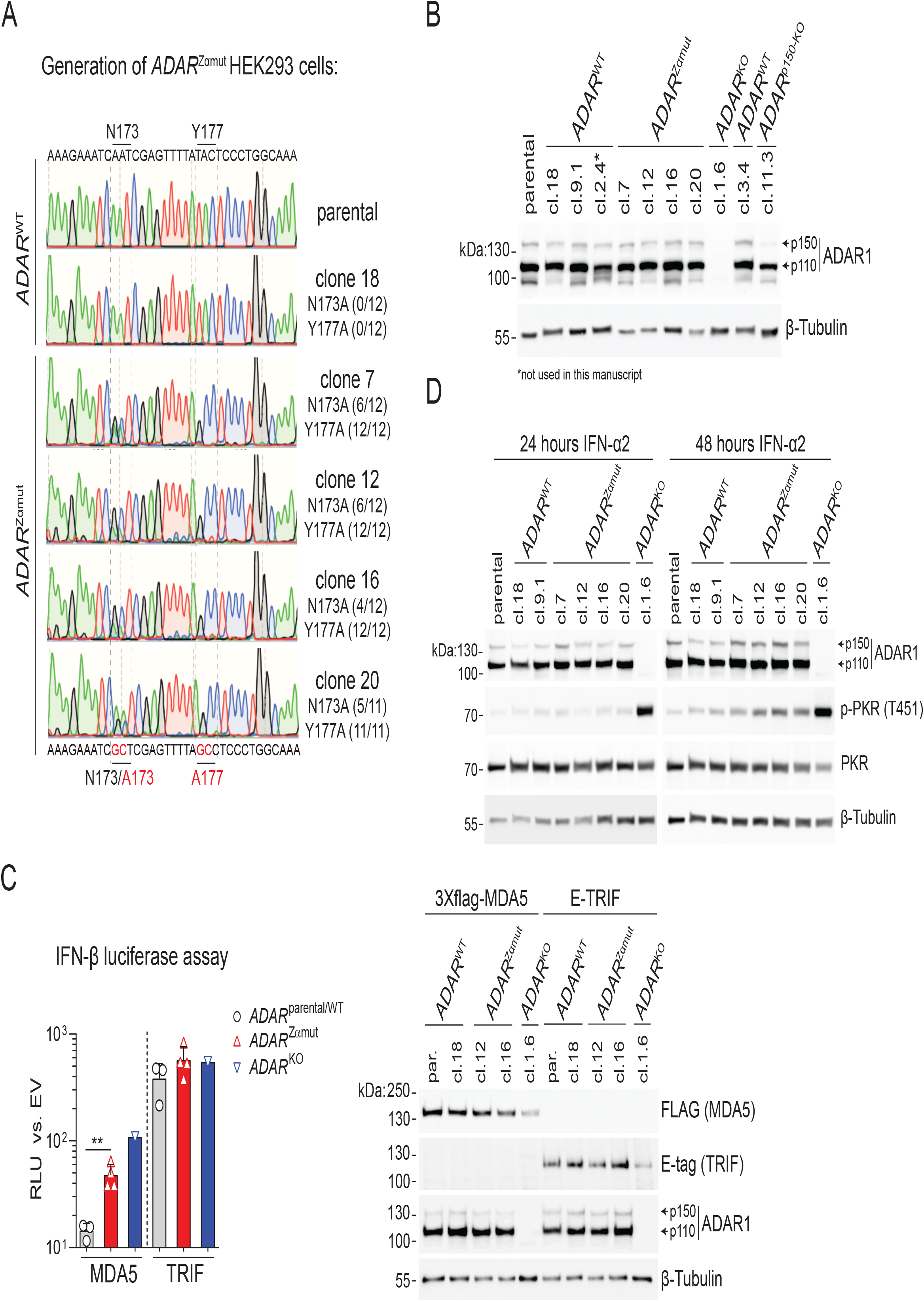
ADAR1 Zα-domain mutation activates MDA5 in human cells. **(A)** Sanger sequencing profiles of the parental HEK293 cells and ADAR1 wild type (*ADAR*^WT^) and Zα-domain mutant (*ADAR*^Zαmut^) HEK293 clones. Ratios indicate the frequency of wild type (N173/Y177) or mutant (A173/A177) alleles determined by Sanger sequencing of 11 or 12 PCR subclones. **(B)** HEK293 clones of the indicated genotypes were stimulated for 24 h with 1000 U/ml IFN-α2. Protein expression of ADAR1 was analysed by Western blotting. Arrows indicate the ADAR1 p110 and p150 isoform. **(C)** Parental HEK293 cells, *ADAR*^WT^, *ADAR*^Zαmut^ or ADAR1-deficient (*ADAR*^KO^) clones were transfected with 50 ng IFN-β firefly luciferase and 20 ng *Renilla* luciferase reporter plasmids, together with FLAG-tagged human MDA5 or E-tagged human TRIF. Luciferase activity was analysed as in Figure 1B. Protein expression of ADAR1, FLAG-tagged MDA5 and E-tagged TRIF was verified by Western blot (right panel). Arrows indicate the ADAR1 p110 and p150 isoform. **, P < 0.01 by unpaired t-test. **(D)** HEK293 clones of the indicated genotypes were stimulated for 24 or 48 h with 1000 U/ml IFN-α2. Protein expression of ADAR1, threonine 451 (T451) phosphorylated PKR (p-PKR) and total PKR was analysed by Western blotting. Arrows indicate the ADAR1 p110 and p150 isoform. Data in (C) and (D) are representative of at least two independent experiments.

**Supplemental Figure 2.**
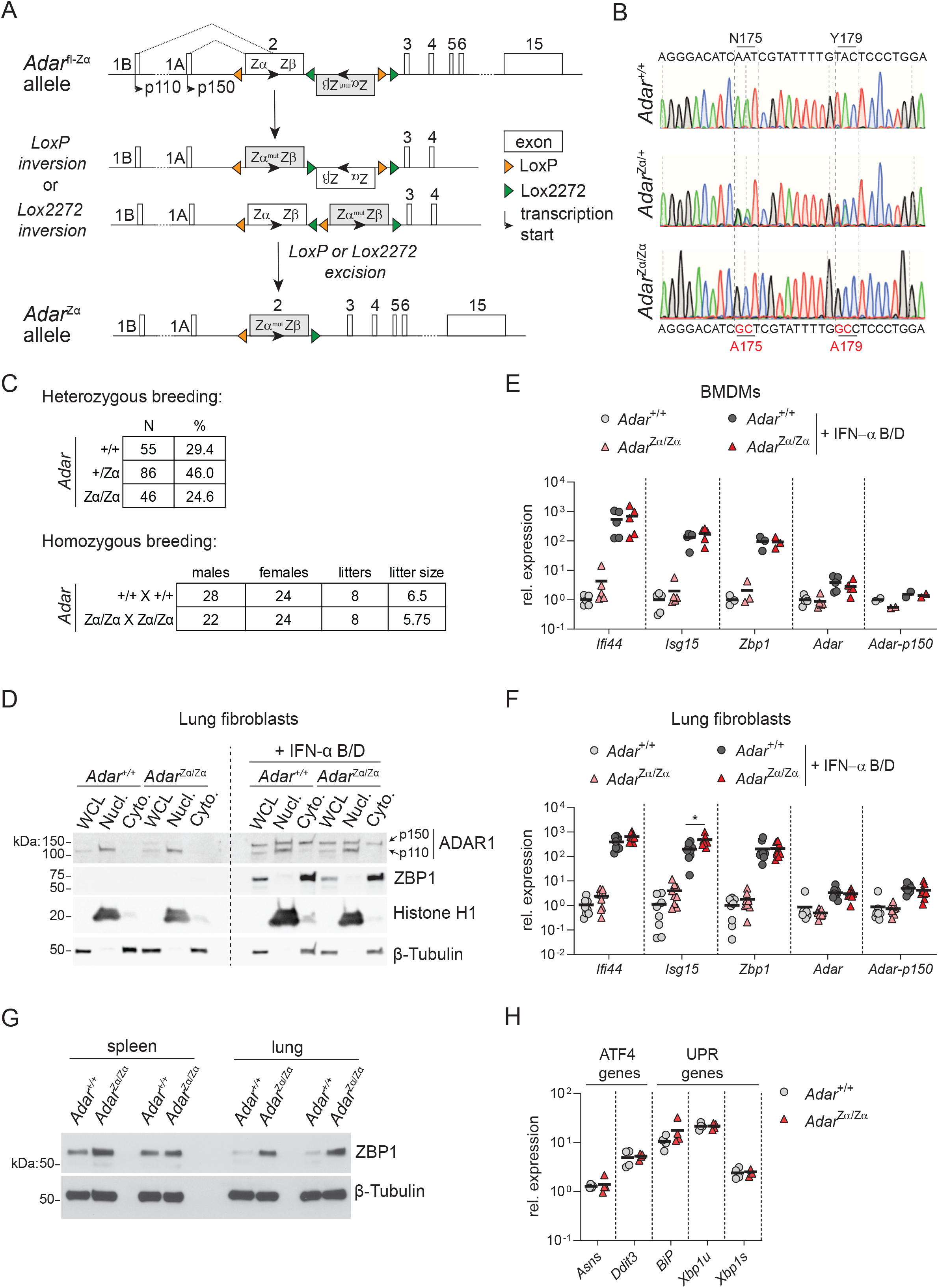
*Adar* Zα-domain mutant mice develop a spontaneous antiviral immune response. **(A)** Schematic overview of the strategy to create the conditional ADAR1 N175A/Y179A Zα-domain knock-in (*Adar*^fl-Zα^) mice. **(B)** Sanger sequencing chromatograms of *Adar*^+/+^, *Adar*^*+/*Zα^ and *Adar*^Zα/Zα^ mice. **(C)** Overview of the offspring obtained from both heterozygous (*Adar*^*+/*Zα^ X Adar^*+/*Zα^) and homozygous (*Adar*^Zα/Zα^ X *Adar*^Zα/Zα^) breeding pairs. **(D)** Primary lung fibroblasts from *Adar*^+/+^ and *Adar*^Zα/Zα^ mice were left untreated or stimulated for 18 hours with 200 U/ml IFN-α B/D. Protein samples were fractioned into whole-cell lysates (WCL), nuclear (Nucl.) or cytosolic (Cyto.) lysates. Protein expression of ADAR1 (isoform p110 and p150), ZBP1, Histone H1 and β-tubulin were analysed by Western blot. **(F)** RT-qPCR analysis of the indicated ISGs in primary lung fibroblasts from *Adar*^+/+^ and *Adar*^Zα/Zα^ mice left untreated or stimulated for 18 hours with 200 U/ml IFN-α B/D. Data are pooled from two independent experiments. The mean relative expression of each gene in untreated *Adar*^+/+^ fibroblasts is set at 1. *, P < 0.05 by unpaired t-test. **(G)** Protein samples were prepared from spleens and lungs of 2 individual *Adar*^+/+^ and Adar^Zα/Zα^ mice. Protein expression of the ISG ZBP1 was analysed by Western blot. **(H)** RT-qPCR analysis of *Asns, Ddit3, BiP* (or *Hspa5*) and *Xbp1* in the unspliced (*Xbp1u*) and spliced (*Xbp1s*) form in the lungs of *Adar*^+/+^ and Adar^Zα/Zα^ mice.

**Supplemental Figure 3.**
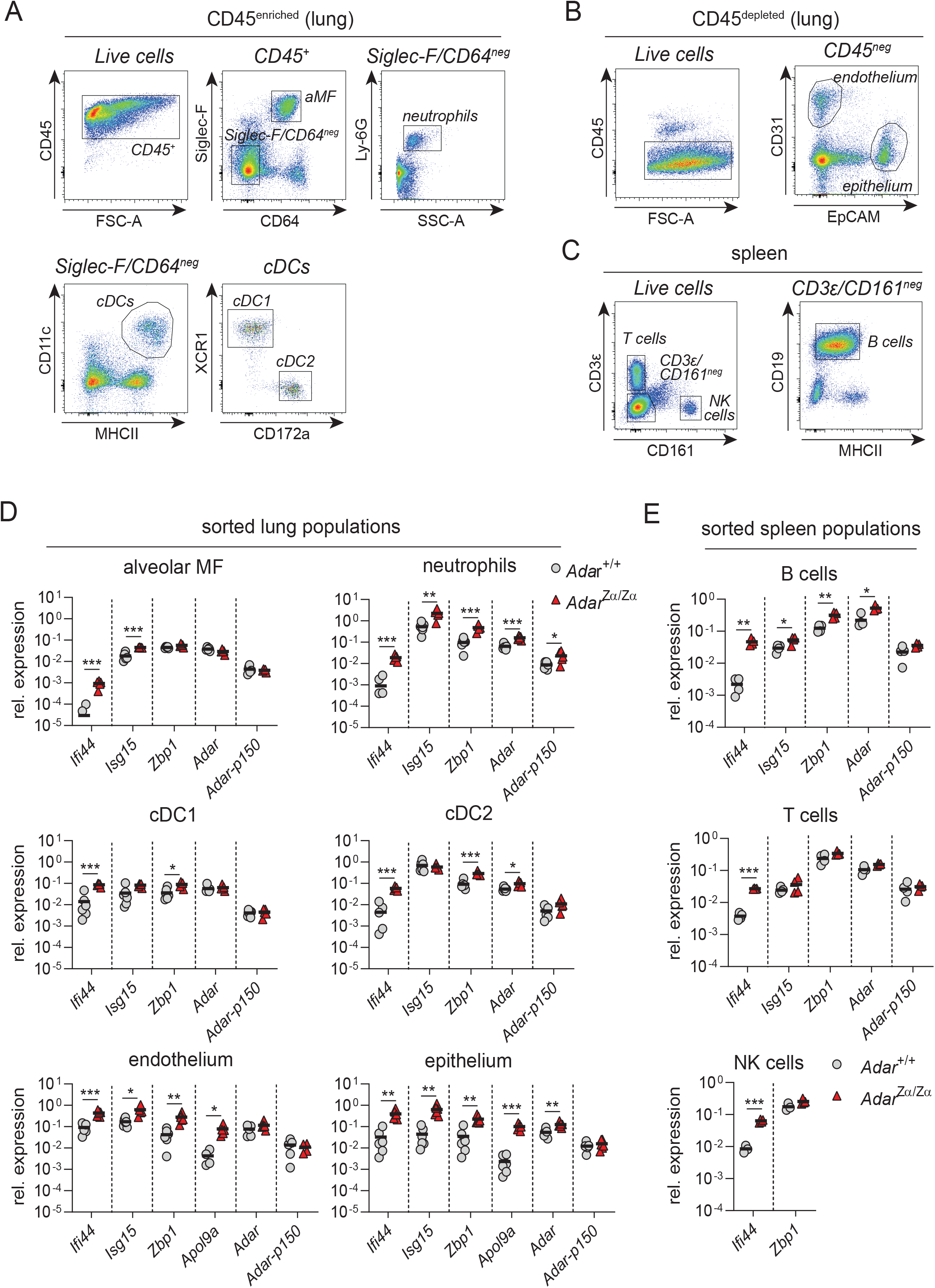
Characterisation of ISG expression in lung and spleen cell types of *Adar*^Zα/Zα^ mice. **(A and B)** Gating strategy for FACS purification of CD45^+^ leukocytes (A) including alveolar macrophages, neutrophils, conventional type 1 and type 2 dendritic cells (cDC1 and cDC2) and CD45^-^ (B) endothelial and epithelial cells from lungs of *Adar*^Zα/Zα^ mice and control littermates. **(C)** Gating strategy FACS purification of B cells, T cells and NK cells from the spleens of *Adar*^+/+^ and *Adar*^Zα/Zα^ mice. **(D and E)** RT-qPCR analysis of the indicated ISGs in the indicated cell populations derived from the lungs (D) and spleens (E) of *Adar*^+/+^ and *Adar*^Zα/Zα^ mice. *, P < 0.05, **, P < 0.01, ***, P < 0.001 by unpaired t-test. Each data point in (D) and (E) represents an individual mouse. Lines indicate the mean. Data in (D) are representative of at least 2 independent experiments.

**Supplemental Figure 4.**
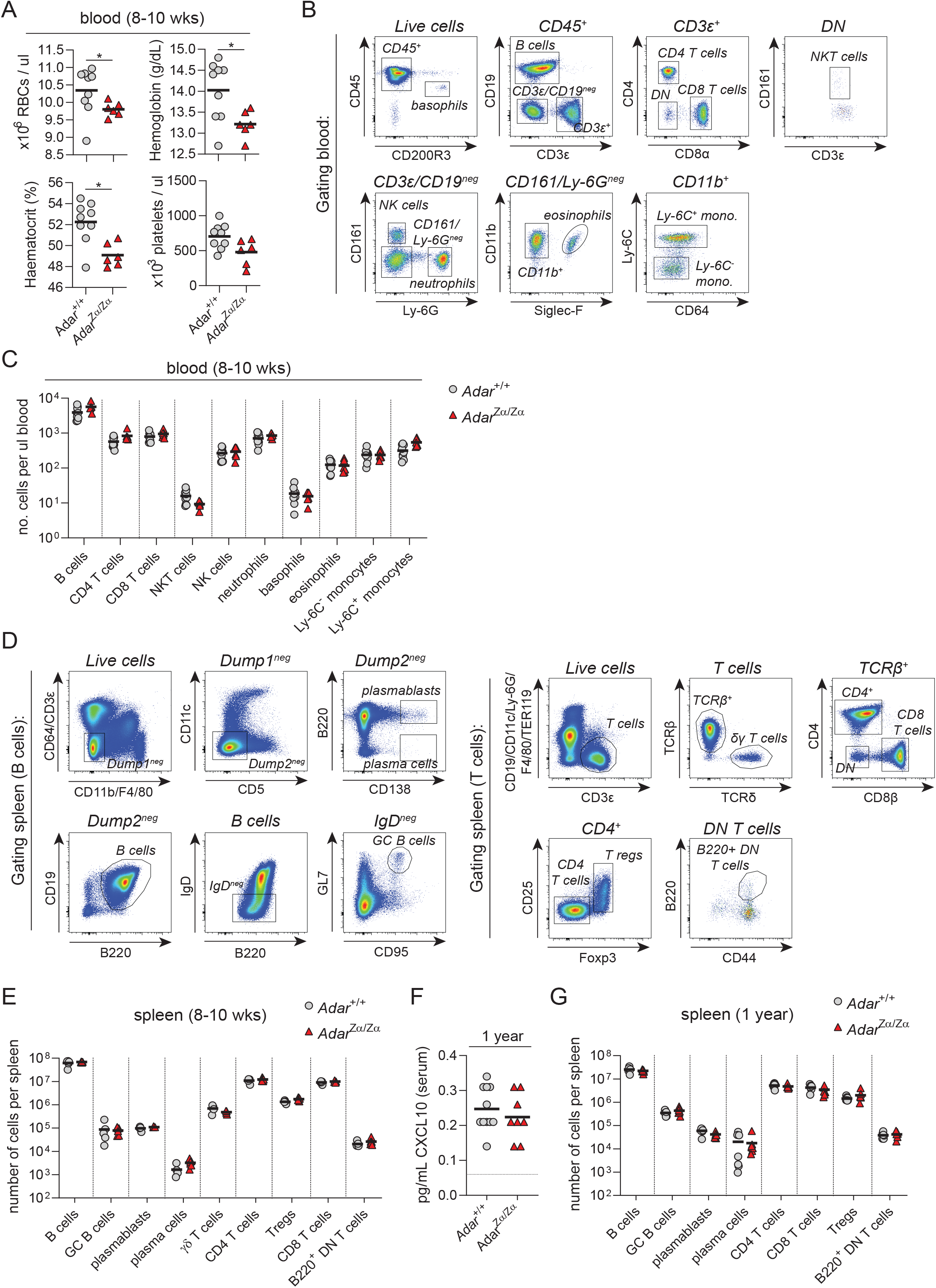
*Adar*^Zα/Zα^ mice maintain normal haematopoiesis and do not develop autoinflammatory disease. **(A)** Peripheral blood from 8-10 week old *Adar*^Zα/Zα^ mice and their wild-type littermates (*Adar*^+/+^) was analysed for total red blood cell and platelets numbers and haematocrit and haemoglobin levels. *, P < 0.05 by Mann–Whitney U test. **(B and C)** Gating strategy (B) used for flow cytometry analysis of circulating lymphocytes (B cells, CD4 and CD8 T cells, and NK and NKT cells) or myeloid cells (neutrophils, basophils, eosinophils, and Ly-6C^-^ and Ly-6C^+^ monocytes) (C). **(D, E and G)** Gating strategy (D) used for flow cytometry analysis of the splenic T and B cell compartment of 8-10 week (E) and 1 year (G) old *Adar*^+/+^ and *Adar*^Zα/Zα^ mice. **(F)** Analysis of CXCL10 protein levels in serum of 1 year old *Adar*^+/+^ and *Adar*^Zα/Zα^ mice by Bio-Plex assay. Each data point in (A), (C), (E-G) represents an individual mouse. Lines indicate the mean.

**Supplemental Figure 5.**
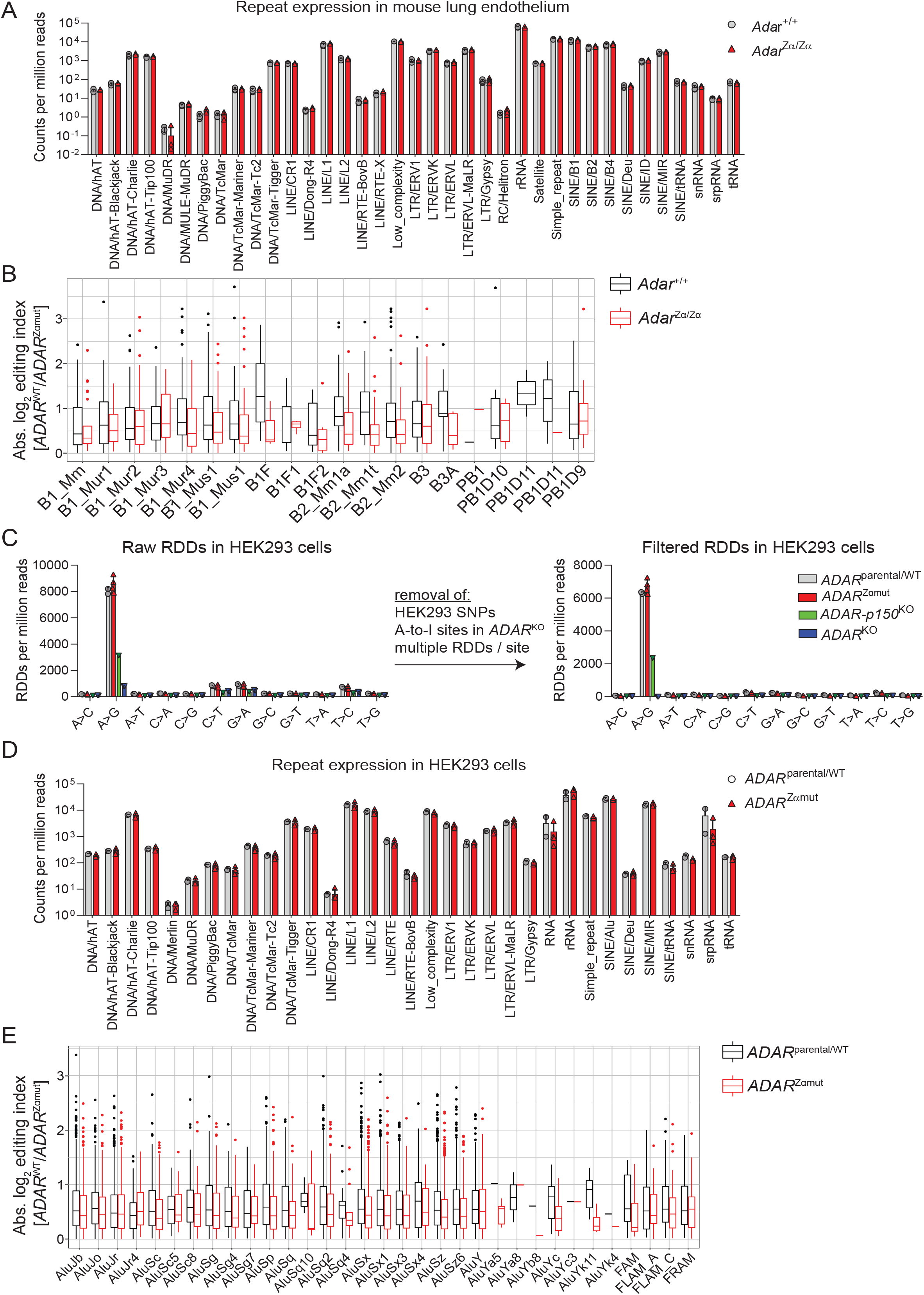
Loss of ADAR1/Z-RNA interaction results in reduced A-to-I editing of SINEs. **(A)** Expression levels in counts per million reads (CPM) of repetitive elements in lung endothelial cells of *Adar*^+/+^ and *Adar*^Zα/Zα^ mice **(B)** box plot showing differential A-to-I editing of members of the mouse SINE B1 and B2 families in lung endothelial cells of *Adar*^+/+^ and *Adar*^Zα/Zα^ mice calculated using the same approach as in Figure 5 (C) and (D). **(C)** Bar charts displaying RNA-DNA differences (RDDs) per million reads in parental HEK293 cells, wild type (*ADAR*^WT^), Zα-domain mutant (*ADAR*^Zαmut^), ADAR1-p150 deficient (*ADAR-p150*^KO^) and ADAR1-deficient (*ADAR*^KO^) HEK293 clones. Left panel: raw RDDs discovered after RDDpred and BAMreadcount. Right panel: refined dataset after removing sites containing multiple RDDs, A>G mismatches discovered in the *ADAR*^KO^ clones and HEK293-specific SNPs. **(D)** Expression levels in counts per million reads (CPM) of repetitive elements in HEK293 cells of the indicated genotype. **(E)** box plot showing differential A-to-I editing of members of the human SINE Alu family in *ADAR* wild type (*ADAR*^parental/WT^) and Zα-domain mutant (*ADAR*^Zαmut^) HEK293 cells calculated using the same approach as in Figure 5 (C) and (D).

